# Canonical Notch2 Signaling Regulates the Development of Iron-recycling Macrophages and Iron Homeostasis

**DOI:** 10.1101/2024.11.13.622461

**Authors:** Frauline Nicole Schroth, Tamar Kapanadze, Stefan Sablotny, Yuangao Xu, Bo Mee Chung, Lena Deuper, Andreas Kispert, Matthias Lochner, Tibor Kempf, Hermann Haller, Kai M. Schmidt-Ott, Jaba Gamrekelashvili, Florian P. Limbourg

## Abstract

Red pulp macrophages (RPM) and bone marrow macrophages (BMM) are iron-recycling cells involved in iron homeostasis and erythropoiesis. Here, we show, using conditional deletion strategies of Notch signaling components, that the development of RPM and BMM is regulated by canonical Notch2 signaling. Loss of functional *Notch2*, or its nuclear mediator *Rbpj*, causes impairment in RPM and BMM and iron overload in the spleen and bone marrow, while prototypic RPM genes required for iron handling are downregulated. This was accompanied by splenic extramedullary hematopoiesis and changes in splenic microarchitecture. Furthermore, early postnatal transfer of bone marrow and fetal liver progenitors rescued the defects in RPM and BMM and iron overload in a *Notch2*-dependent manner, demonstrating the potential to restore defective tissue resident macrophage niches by Notch-competent progenitors. Thus, canonical *Notch2* is required for development and function of iron-recycling macrophages.

## Introduction

Iron demands in mammals are largely supplied through the process of erythrophagocytosis and iron recycling from red blood cells (RBC), which is mainly carried out by iron-recycling macrophages. These are tissue-resident macrophage (TRM) populations comprising splenic red pulp (RP) macrophages (RPM), bone marrow (BM) macrophages (BMM) and liver Kupffer cells (KC)^1^.

The murine spleen (Spl) is organized in discrete microanatomical regions, each performing distinct functions^2^. The unique microanatomy of the Spl promotes the targeted elimination of red blood cells (RBC). This process takes place in the RP, where RPM phagocytose senescent, damaged or infected erythrocytes, hemoglobin or heme complexes, and degrades them. The released iron is then either stored in the cell or shuffled back to the circulation, supplying most of the systemic iron necessary for erythropoiesis^3^. Recently, RPM has been implicated in host defense against pathogens such as *Streptococcus pneumoniae*, underscoring the significance of structure in relation to function^4,5^. The Spl also contains marginal zone (MZ) macrophages (MZM), marginal metallophilic macrophages (MMM) and white pulp (WP) macrophages (WPM)^6^. These macrophages are involved in clearance of blood borne pathogens and control of adaptive immune responses^2^. Splenic CD169^+^ macrophages in the MZ transfer antigens to BATF3-dependent conventional dendritic cells (cDC) that promote the generation of antiviral effector CD8^+^ T cell responses^7^. Clodronate liposome depletion of MZM and MMM in mice causes enhanced spread of *Listeria monocytogenes* to peripheral organs, demonstrating a critical macrophage role in early infection control^8^. MZM depletion also leads to fatal autoimmunity by causing accumulation of exogenously administered apoptotic cells in the WP, further supporting the role of MZM in regulating immune activation and tolerance^9^, while WPM play a crucial role in clearance of apoptotic cells and maintain immune homeostasis within the germinal center^10^.

Like the Spl, the BM hosts distinct populations of macrophages^11^. BMM are specialized resident macrophages that function as iron-rich nurse cells for differentiating erythroblasts. They provide growth factors and iron critically needed for erythrocyte development and hemoglobin synthesis and facilitate erythropoiesis^3,12^. BMM exhibit high levels of proteins involved in iron homeostasis, sharing a similarity to the RPM machinery^13^, and selective deletion of BMM in conditions of erythropoietic stress leads to impaired erythropoietic recovery in mice^12,14,15^.

Previous studies have demonstrated that colony stimulating factor 1 (CSF-1), transcription factor Spi-C (encoded by *Spic*), peroxisome proliferator-activated receptor-γ (PPAR-γ, encoded by *Pparg*), Heme oxygenase 1 (HO-1, encoded by *Hmox1*), BACH1 and Interleukin IL-33 are involved in development, maintenance and function of iron recycling RPM^16–22^. Mice lacking any of these factors exhibit reduced RPM numbers and present an iron overload phenotype in the Spl, emphasizing their essential role in the development and survival of RPM. However, despite recent progress in understanding ontogeny and heterogeneity of mononuclear phagocyte cells including TRM through advanced fate mapping studies, there remains a significant gap in our knowledge of their origin as well as in regulatory factors governing their development and the implications for disease contexts^11,23^.

Notch is a highly conserved signaling pathway that controls lineage commitment and cell development in tissues such as the mononuclear phagocyte system (MPS)^24,25^. For instance, Notch2, and its downstream regulator RBPJ (recombination signal binding protein for immunoglobulin κJ region) have been implicated in the development of specific cDC subsets in the Spl and intestine^26–30^. In the liver, sinusoidal endothelial cells seem to interact with recruited monocytes and drive delta-like 4 (DLL4)-Notch-RBPJ-mediated expression of *Nr1h3* and *Spic* transcription factor genes with subsequent differentiation of monocytes into KC^31,32^. In the context of metabolic dysfunction-associated steatotic liver disease and/or steatohepatitis (MASLD/MASH), deletion of *Rbpj* alters the differentiation trajectory of Ly6C^hi^ monocytes from a pro-pathogenic macrophage phenotype toward a more protective Ly6C^lo^ patrolling monocyte, thereby mitigating some pathological features associated with MASLD/MASH^33^. RBPJ also regulates homeostasis and function of circulating Ly6C^lo^ monocytes by regulating CCR2 expression and controls development of CD16.2^+^ interstitial macrophages in the lung^34^.

Notch signaling interacts with CSF-1, controls macrophage maturation, and promotes arteriogenesis and tissue repair during ischemia^35,36^. Previous work has shown that the DLL1-Notch2 signaling axis promotes the conversion of Ly6C^hi^ monocytes to Ly6C^lo^ patrolling monocytes, elucidating its role in the regulation of the MPS^25,37^. Here, we show using *Cx3cr1^Cre^*-mediated targeting of components of the Notch signaling pathway, a previously unknown requirement for canonical Notch2 signaling in the development of iron-recycling RPM and BMM linked to iron homeostasis and erythropoiesis.

## Results

### Canonical Notch2 signaling controls RPM and BMM development

To examine the role of Notch in the development of MPS, we generated mice with conditional deletion of *Notch2* (*N2*^Δ*Cx3cr1*^) or *Rbpj* (*Rbpj*^Δ*Cx3cr1*^) by crossing *Cx3cr1^Cre^* mice^38^ with *Notch2^lox/lox^* ^39^ or *Rbpj^lox/lox^* animals^40^ and analyzed their offspring. *Cre*-negative littermates served as controls (Ctrl) unless otherwise indicated.

*N2*^Δ*Cx3cr1*^ mice exhibited splenomegaly starting with 4 weeks (wk) of age **(Figure 1A and 1B)** but showed no changes in Spl total cell count and peripheral blood (PB) parameters **(Figure 1C, Figure S1A-1H).** We subjected Ctrl and *N2*^Δ*Cx3cr1*^ Spl cells to flow cytometry and performed unsupervised t-distributed stochastic neighbor embedding (t-SNE) analysis on concatenated LiveCD45^+^Lin^lo/-^CD117^neg^ subsets after exclusion of CD11b/CD11c-double negative cells using a described set of surface markers **(Figure 1D and 1E, Figure S1I)**. This separated several subsets closely resembling Ly6C^hi^ (#1), Ly6C^lo^ (#5) monocytes, CD11c^hi^I-A/I-E^+^ DCs (#8, #9) and F4/80^hi^ RPM (#6, #7) **(Figure 1E and 1F)**. In *N2*^Δ*Cx3cr1*^ mice, the frequency of F4/80^hi^ RPM (#6, #7), CD43^+^Ly6C^lo^ monocytes (#5) and CD11c^hi^CD11b^+^I-A/I-E^+^ DCs (#8) were strongly reduced. At the same time, *N2*^Δ*Cx3cr1*^ mice showed expansion of at least four subsets (#1, #2, #3, #4) of CD11b^+^CD11c^neg^ monocytes which expressed altered levels of Ly6C, CD43 and CD163 **(Figure 1F)**. Thus, *N2*^Δ*Cx3cr1*^ mice show pronounced quantitative and qualitative changes in Spl MPS suggesting that Notch2 regulates development of RPM.

**Figure 1.**
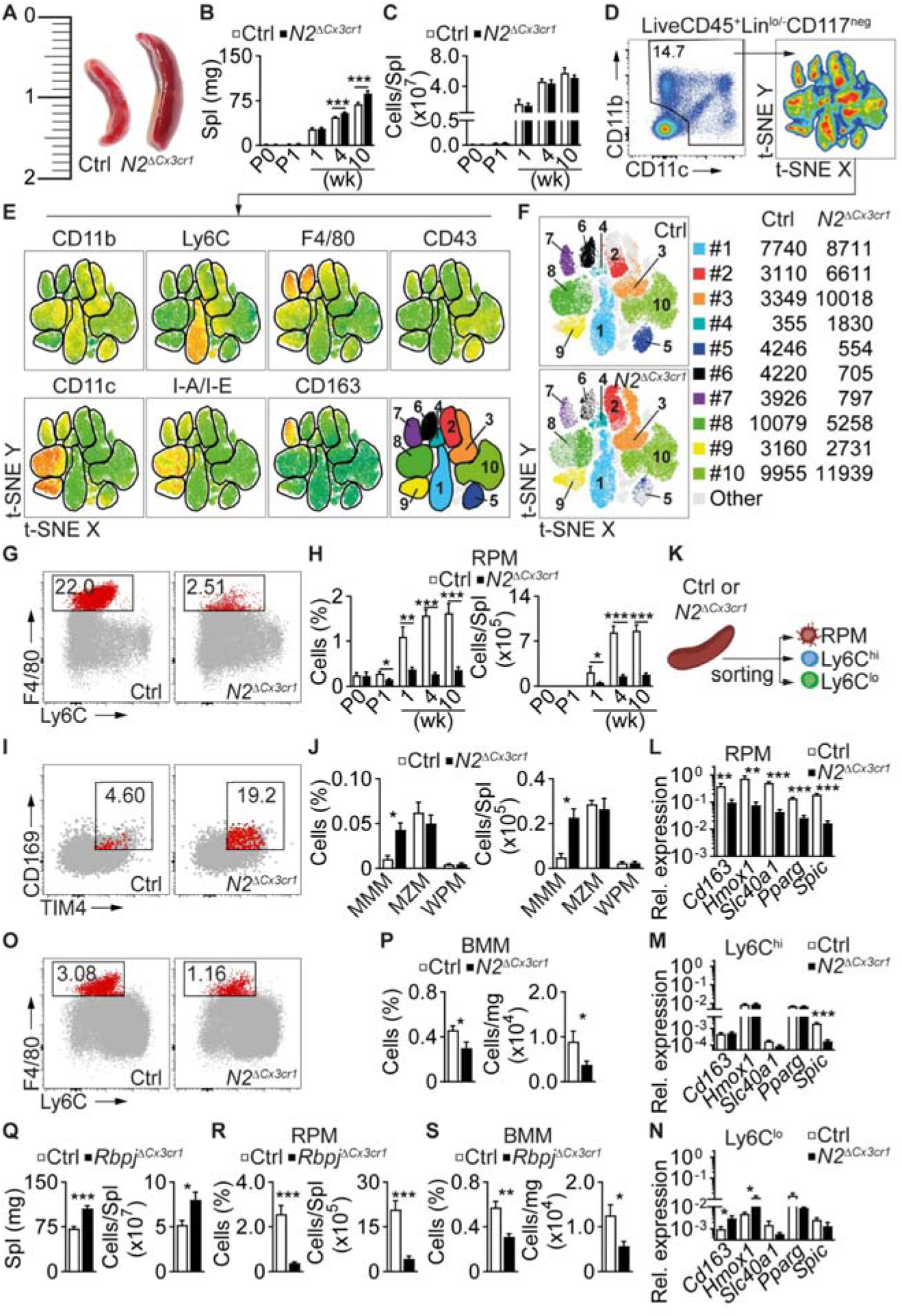
Canonical Notch2 signaling controls RPM and BMM development. (A-C) Splenomegaly in *N2*^Δ*Cx3cr1*^ mice. (A) Spl size, (B) weight and (C) total cell numbers (n = 3-8). (D, E) Gating strategy for t-SNE and definition of splenic phagocytic cells. See also Figure S1I. (D) Unsupervised t-SNE and mapping of MPS subsets on concatenated cells (E) using selected surface markers (n = 6 per group). (F) t-SNE map of control or *N2*^Δ*Cx3cr1*^ Spl (from Figure 1E) showing altered composition of MPS. (G) Representative flow cytometry plots showing Spl RPM. Upstream gating strategy in Figure S1I and S1J. (H) Relative and absolute frequency of Spl RPM at indicated days or weeks after birth (n = 3-8). (I) Representative flow cytometry plot from adult Ctrl or *N2*^Δ*Cx3cr1*^ Spl showing the expansion of CD169^+^ MMM. See also Figure S1K. (J) Relative and absolute frequency of MMM, MZM and WPM in Ctrl or *N2*^Δ*Cx3cr1*^ Spl (n = 6-8). (K) Experimental set-up for cell population sorting for gene expression analysis: RPM (Lin^lo/-^ CD11b^lo/-^Ly6C^lo/-^F4/80^hi^CD115^neg^); Ly6C^hi^ (Lin^lo/-^CD11b^+^Ly6C^hi^F4/80^lo/-^CD115^+^); Ly6C^lo^ (Lin^lo/-^ CD11b^+^Ly6C^lo/-^F4/80^lo/-^CD115^+^). See also Table S1. (L-N) Relative expression of signature genes in RPM and splenic monocytes (n = 6). (O) Representative flow cytometry plots showing BMM in Ctrl or *N2*^Δ*Cx3cr1*^ mice. Upstream gating strategy as in Figure S1I and S1J. (P) Relative frequency and absolute frequency of BMM normalized per mg BM in adult Ctrl or *N2*^Δ*Cx3cr1*^ mice (n = 6-8). (Q-S) Spl weight and total cell number (Q); relative and absolute frequency of RPM (R)- and relative and absolute frequency of BMM (S) in Ctrl or *Rbpj*^Δ*Cx3cr1*^ mice are depicted (n = 6-7). (B, C, H, J, L-N, P-S): Data are mean ± SEM and pooled from, or representative of at least two independent experiments. *P<0.05; **P<0.01; ***P<0.001; (Student’s *t*-test). Lin: Lineage (CD3, CD19, B220, Ly6G, Terr119, NK1.1). See also Figure S1 and Table S1.

Next, we quantified RPM and Ly6C^hi^ monocytes in Spl and BM using a conventional flow cytometry gating strategy **(Figure S1J, Supp. File 1),** comparing Ctrl and *N2*^Δ*Cx3cr1*^ mice from birth (P0, P1) to adulthood (10 wk) **(Figure 1G and 1H).** In Ctrl mice, there was a rapid expansion of RPM from birth to around 1 wk of age with numbers reaching a peak in 4 wk-old animals **(Figure 1G and 1H),** similar to previous reports^21^. This rapid increase likely coincides with the expansion in blood volume and RBC mass during this period^41^. In contrast, RPM numbers in *N2*^Δ*Cx3cr1*^ mice were reduced as early as postnatal day 1 (P1) and remained strongly suppressed into adulthood (10 wk) **(Figure 1G and 1H)**. At the same time, the CD169^+^ MMM population expanded in *N2*^Δ*Cx3cr1*^ mice while the frequency of MZM and WPM was comparable **(Figure 1I and 1J, Figure S1K).** The frequency of splenic Ly6C^hi^ monocytes did not differ between Ctrl and *N2*^Δ*Cx3cr1*^ mice except at 4 wk of age, where there was a marked but transient increase in *N2*^Δ*Cx3cr1*^ animals **(Figure S1L).**

To analyze the expression of RPM signature genes, we sorted RPM and monocytes from Ctrl and *N2*^Δ*Cx3cr1*^ mice and performed quantitative real-time PCR (QRT-PCR) **(Figure 1K)**. The few remaining RPM in *N2*^Δ*Cx3cr1*^ mice exhibit a reduced expression of *Cd163*, *Hmox1, Slc40a1*, *Pparg* and *Spic* **(Figure 1L)**. This coincided with downregulation of *Spic* in Ly6C^hi^ and upregulation of *Cd163* and *Hmox1* in Ly6C^lo^ monocyte subset from *N2*^Δ*Cx3cr1*^ mice, suggesting a potential compensatory response to RPM loss **(Figure 1M and 1N)**. *Spic*-, and *Pparg*-deficient mice^17,21^ show impaired development of BMM. Similarly, the frequency of BMM was also reduced in *N2*^Δ*Cx3cr1*^ mice, suggesting that *Notch2* plays a role in their development **(Figure 1O and 1P)**.

*Cx3cr1* is involved in regulating migration and homing of progenitors or differentiated populations in embryogenesis or after birth^42,43^. To exclude a confounding effect of *Cx3cr1^Cre^* mediated expression of *Cx3cr1*, we analyzed *Cx3cr1^gfp/+^* (heterozygous) and *Cx3cr1^gfp/gfp^* (homozygous) mice lacking one or both alleles of *Cx3cr1*^44,45^. Both strains displayed equal numbers of RPM **(Figure S1M)**, suggesting that *Cx3cr1* loss-of-function does not account for RPM loss.

RBPJ is the key mediator of canonical Notch2 signaling cascade which functions in tandem with the Notch intracellular domain (NICD) to regulate the expression of target genes^46^. Conditional deletion of *Rbpj* in *Rbpj*^Δ*Cx3cr1*^ mice recapitulated the phenotype of *N2*^Δ*Cx3cr1*^ mice. In addition to splenomegaly and increased splenic cell numbers **(Figure 1Q)**, *Rbpj*^Δ*Cx3cr1*^ mice also exhibited substantial loss of RPM and BMM **(Figure 1R and 1S**). Taken together, our data suggests that canonical Notch2 signaling controls RPM and BMM development in Spl and BM, respectively.

### CD163-expressing monocytes expand in *N2*^Δ*Cx3cr1*^ mice

CD163 is a scavenger receptor involved in binding and internalization of haptoglobin-hemoglobin complexes^47^. Our gene expression analysis revealed that *N2*^Δ*Cx3cr1*^ Ly6C^lo^ monocytes upregulate *Cd163* **(Figure 1N)**, while t-SNE analysis showed an expansion of at least four monocyte subpopulations (#1-4) in *N2*^Δ*Cx3cr1*^ mice **(Figure 1F)**. We hypothesize that RPM loss triggers the gradual expansion, from birth, of CD163-expressing monocytes as a compensatory mechanism in *N2*^Δ*Cx3cr1*^ mice. Using a conventional gating strategy built after t-SNE results, we quantified splenic monocyte populations #1, #2, #3 and #4 and CD163-expression over time **(Figure 2A-C)**. Compared to Ctrl mice, *N2*^Δ*Cx3cr1*^ mice showed a gradual and sustained expansion of most monocyte subsets, starting with subset #2 in neonates (1 wk) but ultimately affecting subsets #1 and #3 from the age of 4 wk, while subset #4 was initially decreased but later recovered (**Figure 2D-F**). Simultaneously, the frequency of CD163^+^ cells within monocytes (**Figure 2G-I**), and individual CD163 expression levels (**Fig. 2J-O**), were strongly increased in all subsets in *N2*^Δ*Cx3cr1*^ mice. At the same time, the frequency of CD163^+^ RPM was reduced early on and remained strongly suppressed, while CD163 expression in RPM was initially elevated but later returned to normal levels. Overall, this suggests an early compensatory response to RPM deficiency in mutant mice consisting of CD163 upregulation in RPM and increased monocyte expansion, followed by a late compensatory response consisting of expansion of CD163^+^ monocytes.

**Figure 2.**
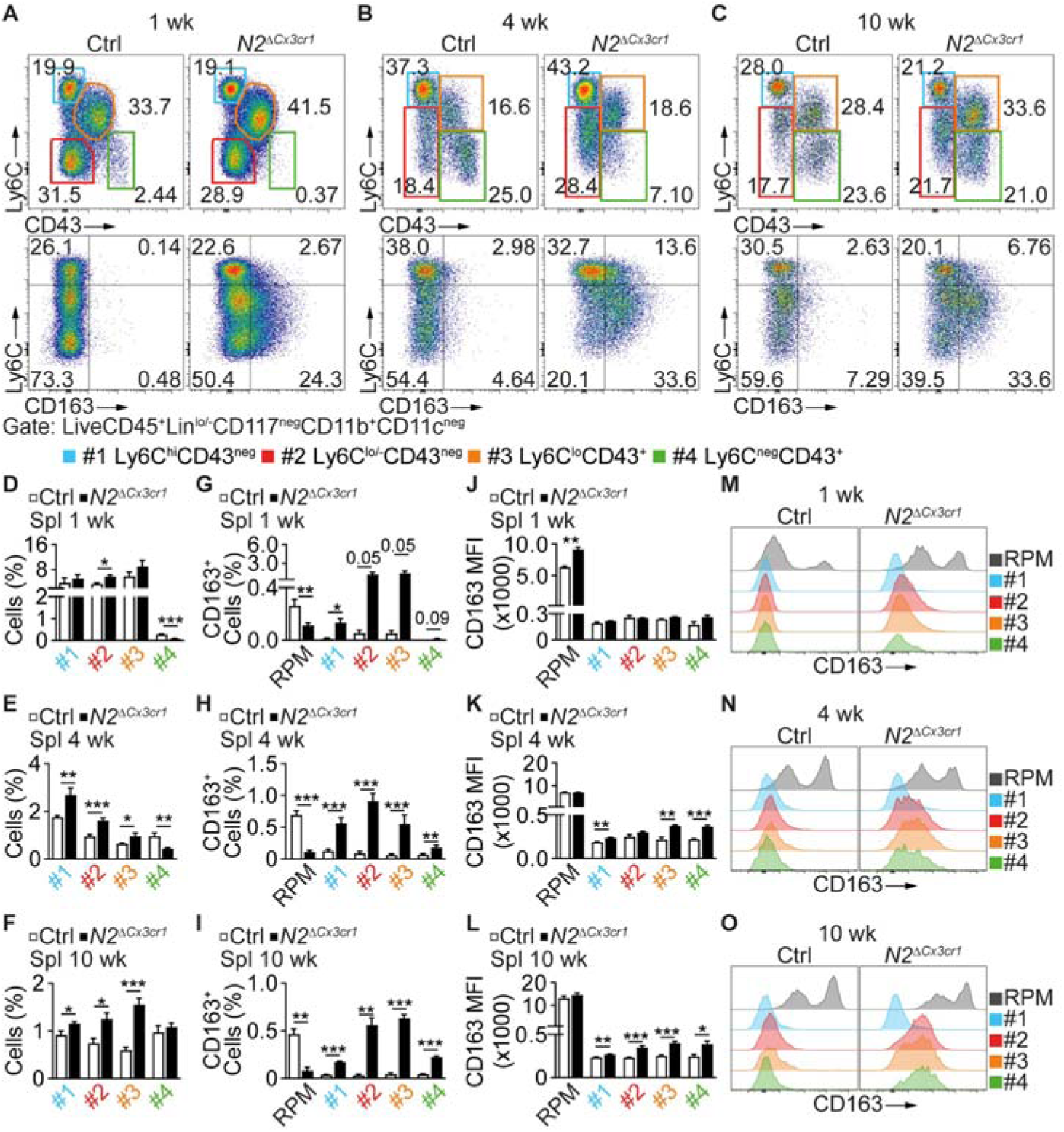
CD163-expressing monocytes expand in *N2*^Δ*Cx3cr1*^ mice. (A-C) Representative flow cytometry plots showing Spl monocyte populations (#1-4) and expression of CD163 in monocytes from 1-, 4- and 10 wk old mice. (D-O) Analysis of monocyte subpopulations in 1-, 4- and 10 wk old mice. (D-F) relative frequency of monocyte populations. (G-I) Relative frequency of CD163^+^ cells within RPM and monocyte populations. (J-L) MFI of CD163 and (M-O) representative histograms showing CD163 expression. (D-L) Data are mean ± SEM and pooled from two or more independent experiments (n = 3-6). *P<0.05; **P<0.01; ***P<0.001; (Student’s *t*-test).

### Impaired splenic microarchitecture and iron homeostasis in canonical *Notch2* signaling-deficient mice

We next investigated the splenic microarchitecture and iron metabolism. Confocal laser scanning microscopy (CLSM) revealed a normal splenic architecture in Ctrl mice, as described by Bellomo et al.^2^. Specifically, there was strong, homogenous F4/80 and CD163 co-staining in the RP, which was separated from the WP by a defined CD169^+^ MZ ring structure, consisting of MMM (**Figure 3A and 3B**). In contrast, in *N2*^Δ*Cx3cr1*^ and *Rbpj*^Δ*Cx3cr1*^ mice, F4/80 staining was reduced and the MZ ring structure was disrupted, with a shift of the CD169 signal to the RP colocalizing with CD163 (**Figure 3A and 3B**). No changes were observed in CD68, which stains RPM and WPM, as well as MARCO and Tim4, which stain subsets of MZM and MMM (**Figure S2A-2D**). Similarly, F4/80, CD169 and Tim4 staining, as well as splenic microarchitecture remained unchanged in *Cx3cr1^gfp/+^* and *Cx3cr1^gfp/gfp^*mice (**Figure S2E and 2F**), confirming data obtained by flow cytometry (**Figure S1M**). Collectively, these data show that the deletion of *Notch2* or *Rbpj* significantly alters the splenic microarchitecture due to changes in RPM and MMM in the splenic niche.

**Figure 3.**
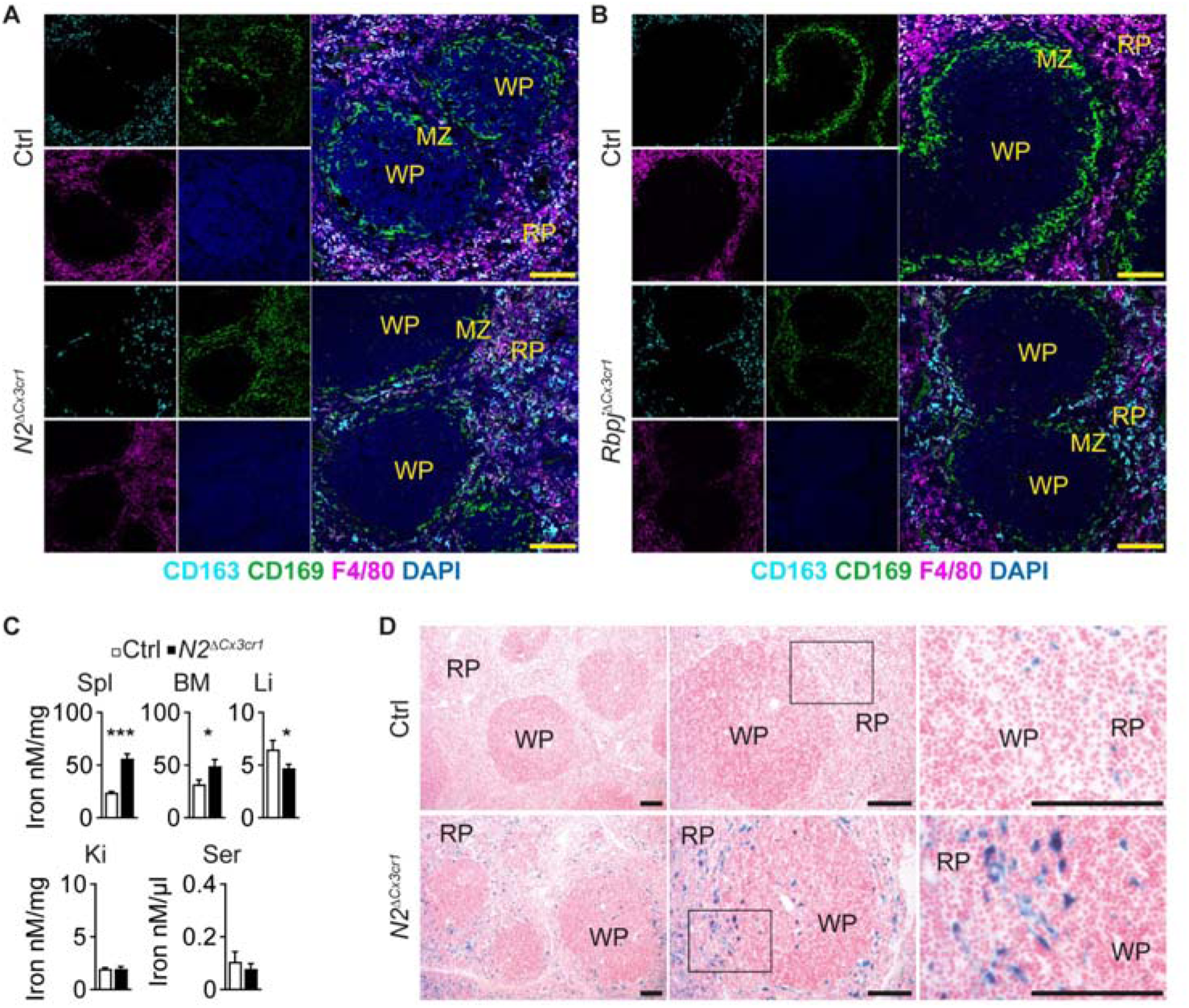
Impaired splenic microarchitecture and iron homeostasis in canonical *Notch2* signaling-deficient mice. (A, B) CLSM images of Spl from 10 wk old (A) Ctrl or *N2*^Δ*Cx3cr1*^- or (B) Ctrl or *Rbpj*^Δ*Cx3cr1*^ mice. (C) Iron quantification in Spl, BM, Liver (Li), Kidney (Ki) and Serum (Ser) of mice. Data are mean ± SEM and pooled from two or more independent experiments (n = 3-10). *P<0.05; **P<0.01; ***P<0.001; (Student’s *t*-test). (D) Perl’s Prussian blue staining showing iron deposits in 10 wk old Spl of Ctrl and *N2*^Δ*Cx3cr1*^ mice. (A, B, D) Images are representative of two or more independent experiments. Scale bar 100 µm. See also Figure S2 and S3.

Next, we tested whether changes in the RPM pool result in impaired iron recycling in *N2*^Δ*Cx3cr1*^ mice. Despite normal PB parameters (**Figure S1A-1H**), there was iron accumulation in Spl and BM, but not in kidney or serum, and a reduction of iron content in the liver in *N2*^Δ*Cx3cr1*^ mice (**Figure 3C, Figure S3**). Perl’s Prussian Blue staining confirmed visible and focal iron deposition in the RP of *N2*^Δ*Cx3cr1*^ Spl sections (**Figure 3D**).

### Deletion of *Notch2* does not affect RPM proliferation or cell death

To investigate whether proliferation differences could account for the early onset impairment in RPM development, we performed Ki-67 staining. *In situ* analysis by CLSM showed prominent Ki-67 staining in the RP of both Ctrl and *N2*^Δ*Cx3cr1*^ mice at the age of 2 wk, which was absent in 10 wk-old mice, indicating transient proliferation preceding full maturation (**Figure S4A and 4B**). The majority of these proliferating cells were located in the RP with a few Ki-67^+^ cells in the WP (**Figure S4A**). We found bright F4/80^+^ cells that partially colocalized with Ki-67 in the RP, and with concentric CD169^+^ MMM rings in young Ctrl mice, suggestive of the proliferative capacity of RPM during the onset of normal splenic architecture formation (**Figure S4A and 4B**). In contrast, young *N2*^Δ*Cx3cr1*^ Spl already showed the evident loss of RPM and aberrant CD169 staining in the RP and although Ki-67 was present, these cells were not positive for F4/80 (**Figure S4A and 4B**). BrdU incorporation in RPM was similar in both strains (**Figure S4C**), suggesting that deletion of *Notch2* does not affect proliferation capacity of RPM. We also analyzed cell death by Annexin V (AnnV) and propidium iodide (PI) staining of Spl cells. There was no significant difference in the number of apoptotic or necrotic cells within the RPM population from Ctrl and *N2*^Δ*Cx3cr1*^ mice (**Figure S4D**). Thus, deletion of *Notch2* affects neither death nor proliferation ability of RPM.

### Reconstitution of the splenic niche with BM monocyte-derived RPM

To test whether Notch2 controls cell differentiation or cell commitment to an RPM fate, we performed several adoptive transfer studies. First, we transferred CD45.1^+^ whole BM or whole Spl cells into CD45.2^+^ *N2*^Δ*Cx3cr1*^ neonates and analyzed the CD45.1^+^ donor population 8-14 weeks later (**Figure 4A, Figure S5A**). BM, but not Spl, cells gave rise to RPM which reconstituted the defective niche and rescued the *N2*^Δ*Cx3cr1*^ phenotype, while donor Spl cells failed to do so (**Figure 4B-4D and Figure S5B and S5C**). Neither BM nor Spl cell transfer rescued BMM (**Figure 4E and 4F**) despite the presence of a distinct donor-derived population (∼30%) in recipients of BM (**Figure 4F**), nor did it change the total number of BM cells in recipients (**Figure S5C**).

**Figure 4.**
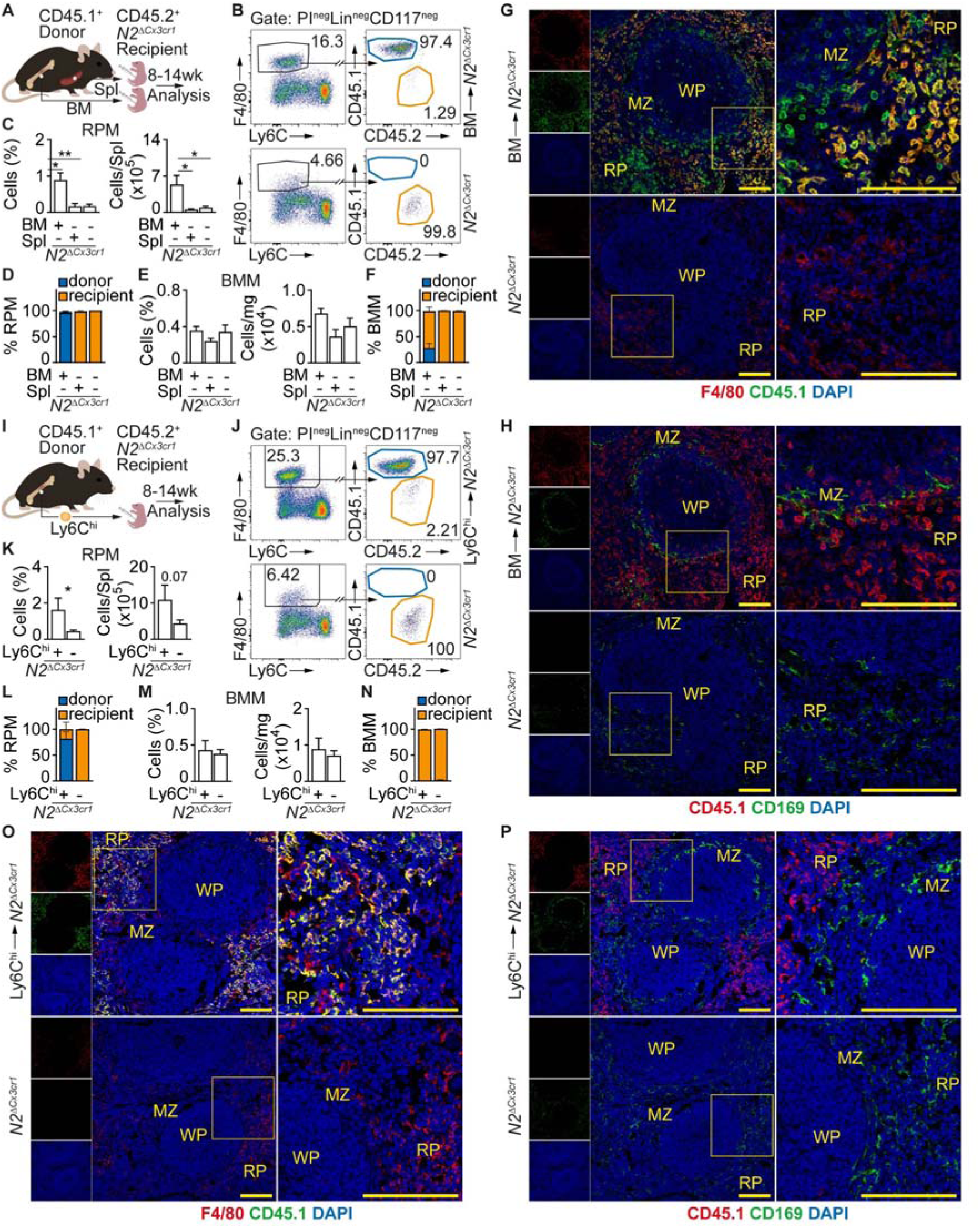
Reconstitution of the splenic niche with BM monocyte-derived RPM. (A) Experimental scheme of reconstitution studies. (B) Representative flow cytometry plot depicting donor-derived RPM in *N2*^Δ*Cx3cr1*^ recipient Spl. See also Figure S5A. (C) Relative and absolute frequency of RPM in recipient mice. (D) Frequency of donor- and recipient-derived RPM in whole RPM pool. (E) Relative and absolute frequency of BMM calculated per mg BM in recipient mice. (F) Frequency of donor- and recipient-derived BMM in whole BMM pool of recipients reconstituted with BM or Spl cells. (G, H) CLSM images of Spl from BM-reconstituted-, or non-reconstituted *N2*^Δ*Cx3cr1*^ mice. Images are from the same sample but depicted separately for simplicity. (I) Experimental scheme of reconstitution studies using BM Ly6C^hi^ monocytes. (J) Representative flow cytometry plot depicting donor-derived RPM after reconstitution. (K) Relative and absolute frequency of RPM in recipient mice. (L) Frequency of donor- and recipient-derived RPM in whole RPM pool. (M) Relative and absolute frequency of BMM in recipient mice. (N) Frequency of donor- and recipient-derived BMM in whole BMM pool. (O-P) Spl CLSM images from recipient *N2*^Δ*Cx3cr1*^ mice reconstituted or not with BM Ly6C^hi^ monocytes. Images are from the same sample but depicted separately for simplicity. (C-F, K-N) Data are mean ± SEM and pooled from two or more experiments (n = 3-6). *P<0.05; **P<0.01; ***P<0.001; (Student’s *t*-test). (G, H, O, P) Representative of two or more experiments. Scale bar 100 µm. See also Figure S5.

CLSM revealed a strong F4/80 signal in the RP and defined CD169^+^ ring of MMM in the Spl of BM-reconstituted mice (**Figure 4G and 4H, Figure S5D**), phenocopying the normal splenic architecture seen in Ctrls (**Figure 3A and 3B**). The F4/80^+^ RPM costained with CD45.1, confirming their donor origin. This was not the case for CD169, which did not colocalize with CD45.1 (**Figure 4H**). In line with these data, we saw little or no accumulation of iron in BM-reconstituted *N2*^Δ*Cx3cr1*^ Spl, supporting a phenotype rescue (**Figure S5E andS5F**).

We next hypothesized that BM Ly6C^hi^ monocytes are capable of differentiating into RPM in the defective splenic niche. To test this we transferred sorted CD45.1^+^ BM Ly6C^hi^ monocytes into CD45.2^+^ *N2*^Δ*Cx3cr1*^ newborns and analyzed their fate after 8-14 weeks (**Figure 4I, Figure S5A**). BM Ly6C^hi^ monocytes reconstituted the defective niche and rescued RPM numbers compared to non-reconstituted or BM reconstituted *N2*^Δ*Cx3cr1*^ mice (**Figure 4J-4L**, **C**). As with whole BM reconstitution, RPM were mostly (∼75%) donor-derived (**Figure 4L**), while the frequency of Spl CD45^+^ cells remained unchanged (**Figure S3G**). Transfer of Ly6C^hi^ monocytes influenced neither BMM nor total BM cells (**Figure 4M, Figure S5H**) and the BMM pool present in reconstituted *N2*^Δ*Cx3cr1*^ mice was of recipient origin (**Figure 4N**). Again, characteristic F4/80^+^ RPM in the RP and a defined CD169^+^ MMM rings were present in recipients of CD45.1^+^ Ly6C^hi^ monocytes but not in *N2*^Δ*Cx3cr1*^ animals without reconstitution (**Figure 4O and 4P, Figure S5I**). F4/80 colocalized with CD45.1, revealing the donor origin of RPM (**Figure 4O**), but the CD169^+^ MMM ring structure were mainly CD45.1 negative (**Figure 4P**). Despite successful repopulation with BM Ly6C^hi^ monocytes, the Spl of reconstituted *N2*^Δ*Cx3cr1*^ mice showed iron deposits in the RP (**Figure S5J-S5K**). Thus, BM Ly6C^hi^ monocytes do not fully functionally rescue the iron accumulation phenotype seen in the *N2*^Δ*Cx3cr1*^ mice.

### Development of RPM and BMM requires intrinsic Notch2 signaling

To directly address the requirement of intrinsic *Notch2* for RPM and BMM development, we generated *N2*^Δ*Cx3cr1*^ *mG* (KO *mG*) and Ctrl *mG* strains by crossing *ROSA26-mT/mG* mice^48^ with *Cx3cr1^Cre^ Notch2^lox/lox^ (N2*^Δ*Cx3cr1*^) or *Cx3cr1^Cre^ Notch2^+/+^ (*Ctrl) animals, respectively. This targeting strategy leads to the simultaneous deletion of *Notch2* and expression of *mG* in *Cx3cr1*-expressing cells and their progeny, which can be used for repopulation and subsequent cell fate studies specifically addressing the role of Notch2.

We transferred BM cells from Ctrl *mG* or KO *mG* mice into newborn *N2*^Δ*Cx3cr1*^ animals and analyzed the recipients 8-14 wk later (**Figure 5A**, **Figure S6A**). Ctrl *mG* BM, but not KO *mG* BM restored RPM, which were donor-derived (**Figure 5B-5D**), and prevented splenomegaly without altering total splenic CD45^+^ cells in recipients (**Figure S6B and S6C**). Furthermore, the transfer of Ctrl *mG* BM cells in *N2*^Δ*Cx3cr1*^ mice also rescued BMM, which were mostly (∼80%) of donor origin but the total number of BM cells remained unchanged in recipient mice (**Figure 5E and 5F, Figure S6C**).

**Figure 5.**
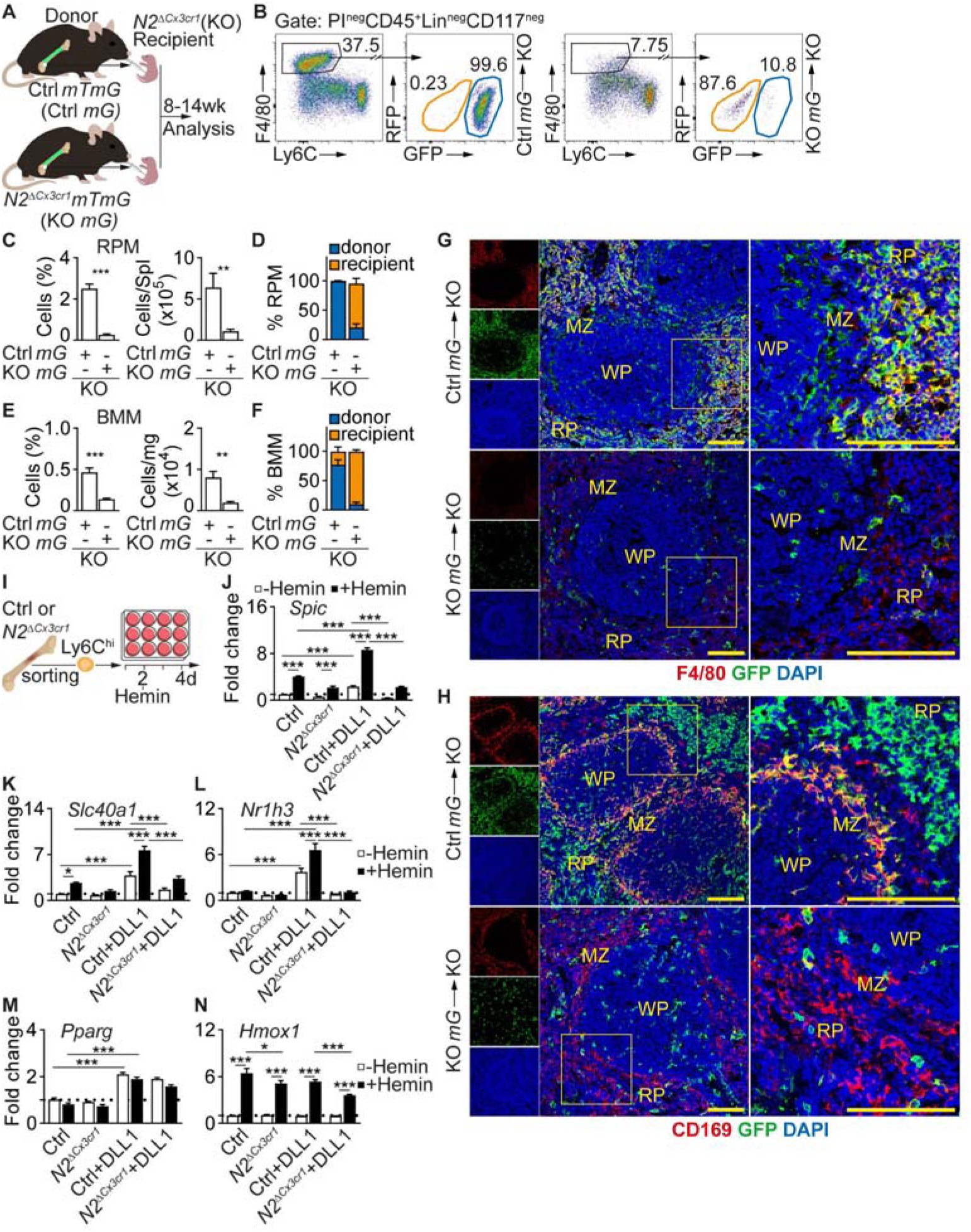
Development of RPM and BMM requires intrinsic Notch2 signaling. (A) Experimental scheme of reconstitution study. (B) Representative flow cytometry plot depicting donor-derived RPM in *N2*^Δ*Cx3cr1*^ recipients. S Figure S6A. (C) Relative and absolute frequency of RPM in recipient mice. (D) Frequency of donor- and recipient-derived RPM in whole RPM pool. (E) Relative and absolute frequency of BMM calculated per mg BM in recipient mice. (F) Frequency of donor- and recipient-derived BMM in whole BMM pool. (G, H) CLSM images of Spl from Ctrl *mG* or KO *mG* BM-reconstituted *N2*^Δ*Cx3cr1*^ mice. Representative of two or more experiments. Scale bar 100 µm. (I) Experimental scheme: Sorted BM Ly6C^hi^ monocytes cultured *in vitro* on immobilized DLL1 ligand and in the presence of CSF-1 and Hemin and differentiated into RPM-like macrophages. (J-N) Expression of RPM signature genes in RPM-like macrophages generated from Ctrl or *N2*^Δ*Cx3cr1*^ Ly6C^hi^ monocytes *in vitro* (n = 6-8). Bar graphs are mean ± SEM. *P<0.05; **P<0.01; ***P<0.001; Two way ANOVA with Bonferroni’s multiple comparison test. (C-F) Data are mean ± SEM and pooled from two or more experiments (n = 6). *P<0.05; **P<0.01; ***P<0.001; (Student’s *t*-test). See also Figure S6.

By CLSM, mice reconstituted with Ctrl *mG* BM showed F4/80^+^ RPM in the RP and CD169^+^ MMM rings in MZ that stained positive for GFP, revealing donor origin (**Figure 5G and 5H**). In contrast, KO *mG* BM-reconstituted mice contained few F4/80^+^ cells and lacked the defined CD169^+^ MMM rings (**Figure 5G and 5H**). Furthermore, Ctrl *mG* BM-reconstituted animals exhibited lower iron levels and only minimal iron deposits in the Spl, again suggesting a rescued phenotype (**Figure S6D andS6E**), while more and large iron deposits were present in the RP of KO *mG* BM-reconstituted mice (**Figure S6D**).

Previous studies have demonstrated that fetal liver (FL) cells can colonize empty niches and develop into functional tissue resident macrophages including RPM^21,49^. In this context, we studied whether *Notch2* also controls the colonization of Spl niche by FL cells and their subsequent development into RPM. We transferred E14.5 Ctrl *mG*-, and KO *mG* FL cells into *N2*^Δ*Cx3cr1*^ neonates and analyzed recipients at 8-14 wk (**Figure S7A**). Ctrl *mG* FL cells restored RPM and BMM pools with cells of donor origin (**Figure S7B-S7F**) and rescued the splenomegaly phenotype (**Figure S7G and S7H**), while the few remaining RPM and BMM in KO *mG* FL-reconstituted mice were recipient-derived cells (**Figure S7D and S7F**). *In situ* analysis revealed that Ctrl *mG* FL-reconstitution, but not KO *mG* FL-reconstitution, also restored RPM and MMM subsets and splenic microarchitecture (**Figure S7I and S7J**) in recipients similar to Ctrl- or KO *mG* BM-reconstitution (**Figure 5G and 5H**). As expected, Ctrl *mG* FL-reconstituted Spl showed reduced iron deposits in contrast to KO *mG* FL-reconstituted mice, which had abundant diffuse iron inserts in the RP (**Figure S7K and S7L**). Thus, Notch2 is unequivocally required for the development of RPM and BMM from both BM or FL progenitors and for formation of the normal splenic microarchitecture.

### DLL1 - Notch2 regulates heme-induced RPM signature genes *in vitro*

Heme functions as a physiological cue to promote monocyte differentiation into iron-recycling macrophages^17^. To test whether Notch regulates RPM signature genes in response to heme in a defined *in vitro* system, we cultured Ctrl or *N2*^Δ*Cx3cr1*^ BM Ly6C^hi^ monocytes on immobilized recombinant Notch ligand DLL1 in the presence of CSF-1 and hemin and analyzed the resultant macrophages for the expression of signature RPM genes (*Spic*, *Slc40a1*, *Nr1h3*, *Pparg*, *Hmox1*) (**Figure 5I**).

Hemin and DLL1 upregulated *Spic and Slc40a1* (encoding ferroportin) individually and in synergy, which was mediated by Notch2, while *Nr1h3* (encoding LXR-alpha) was strictly dependent on DLL1-Notch2 (**Figure 5J-5L**). *Pparg* was expressed in cells cultured on DLL1, but was not dependent of *Notch2* or hemin (**Figure 5M**). *Hmox1,* in contrast, was strictly hemin-dependent and not regulated by DLL1-Notch2 (**Figure 5N**). These data indicate that DLL1-Notch2 signaling promotes expression of RPM signature genes in conjunction with hemin and that it is required for expression of *Spic, Slc40a1* and *Nr1h3* in RPM development.

### Enhanced extramedullary erythropoiesis in *N2***^Δ^***^Cx3cr1^* mice

During conditions of erythropoietic stress, erythropoiesis shifts to extramedullary sites such as the Spl, thus expanding production of reticulocytes and mature erythrocytes^50^. Analysis of erythroid progenitors^51^ (**Figure S8A and S8B**) revealed that *N2*^Δ*Cx3cr1*^ mice exhibit decreased CD45^neg^ early erythroid progenitors ProE in BM, but increased numbers of erythroid progenitors ProE, EryA and EryB in Spl and minor changes in liver (**Figure 6A-6E**). The mouse Spl sustains an active erythropoietic activity from birth that gradually ceases around 7 weeks of age when the BM establishes itself as the main erythropoietic organ^41^. Independent of genotype, we saw an abundance of proliferating CD71^+^ erythropoietic cells in 2 wk-old mice (**Figure 6F-6G**). In Ctrl mice, these cells formed clusters of isolated erythropoietic islands surrounded by F4/80^+^ RPM, but in *N2*^Δ*Cx3cr1*^ mice CD71^+^ cells were distributed throughout the RP. In 10 wk old mice proliferation was absent regardless of genotype, but in contrast to Ctrl mice, where CD71^+^ cells were absent, *N2*^Δ*Cx3cr1*^ mice maintained abundant clusters of erythropoietic islands (**Figure 6H-6I**). Additional staining revealed smaller, non-nucleated CD71^+^Ter119^+^ double-positive early erythrocytes in proximity to the islands (**Figure 6J**). Furthermore, in *N2*^Δ*Cx3cr1*^ mice CD71^+^ erythropoietic islands were surrounded by CD169^+^ cells, while in Ctrl mice, Ter119^+^ erythrocytes were scattered in between F4/80^+^ RPM in the RP (**Figure 6J, Figure S8C-S8E**). Interestingly, despite the rescue of RPM in *N2*^Δ*Cx3cr1*^ mice after repopulation with the BM or FL cells (**Figure 5, Figure S7**), CD71^+^ cell islands remain in the Spl (**Figure 6K and Figure S8F**). Together, these findings demonstrate loss of RPM and BMM with iron overload and persistence of extramedullary erythropoiesis in mature mice in the absence of canonical Notch2 signaling. This suggests that extramedullary erythropoiesis is a compensatory mechanism for ineffective BM erythropoiesis in mature mice, possible secondary to loss of BMM.

**Figure 6.**
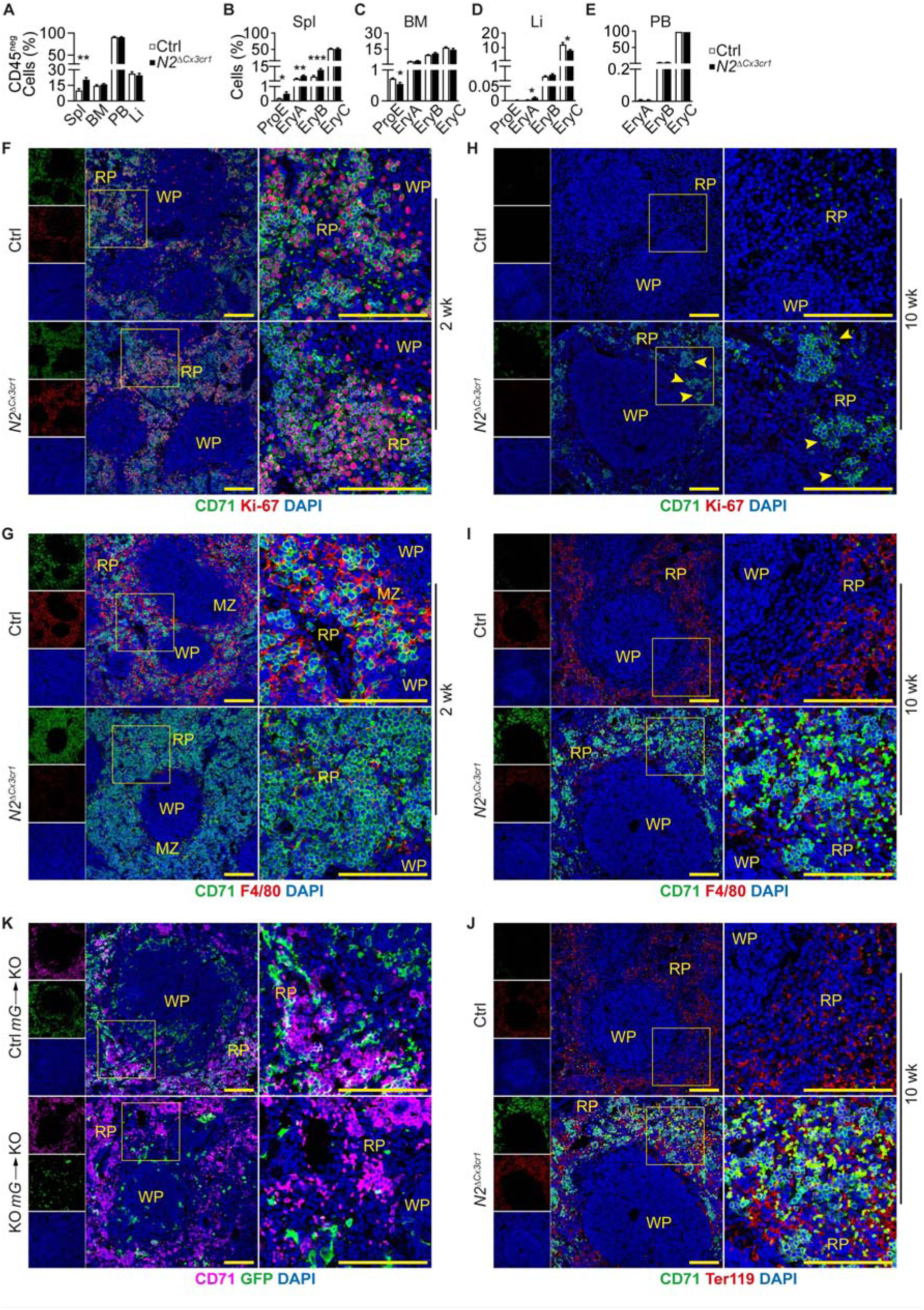
Enhanced extramedullary erythropoiesis in *N2*^Δ*Cx3cr1*^ mice. (A-E) Relative frequency of CD45^neg^ cells (A), erythroid progenitors and mature erythrocytes in the Spl, BM, Liver (Li) and PB (B-E). Bar graphs depict mean ± SEM (n = 8-9). *P<0.05; **P<0.01; ***P<0.001; (Student’s *t*-test). (F-J) Spl CLSM images from 2 wk (F,G), or 10 wk (H-J) old Ctrl or *N2*^Δ*Cx3cr1*^ mice. (H) Arrowheads depict erythropoietic clusters. (F,G) Images are from the same sample but depicted separately for simplicity. (I,J) Images are from the same sample but shown separately for simplicity. See also Figure S8E. (K) CLSM images of *N2*^Δ*Cx3cr1*^ Spl reconstituted with Ctrl *mG* or KO *mG* BM cells. (F-K) Images are representative of at least two independent experiments. Scale bar 100 µm. See also Figure S8 and Table S1.

### *Notch2*-deficiency confers improved adaptation to PHZ-induced hemolytic anemia

To test whether *Notch2* deletion alters the response to anemia, we induced hemolysis through phenylhydrazine (PHZ) administration and analyzed PB and organ parameters (**Figure S9A**). Following PHZ, hemoglobin (HGB) levels decreased significantly faster and more pronounced in Ctrl than in *N2*^Δ*Cx3cr1*^ mice in the early response period, while recovery of hematocrit (HCT) and RBC counts was weaker at d5 (**Figure 7A-7D**, **Figure S9C-S9N**). Mice also developed splenomegaly, which at its peak was not different anymore between genotypes (**Figure S9B**). This observation indicates better adaptation to hemolytic stress in *Notch2*-deficient mice in the initial response of anemia.

**Figure 7.**
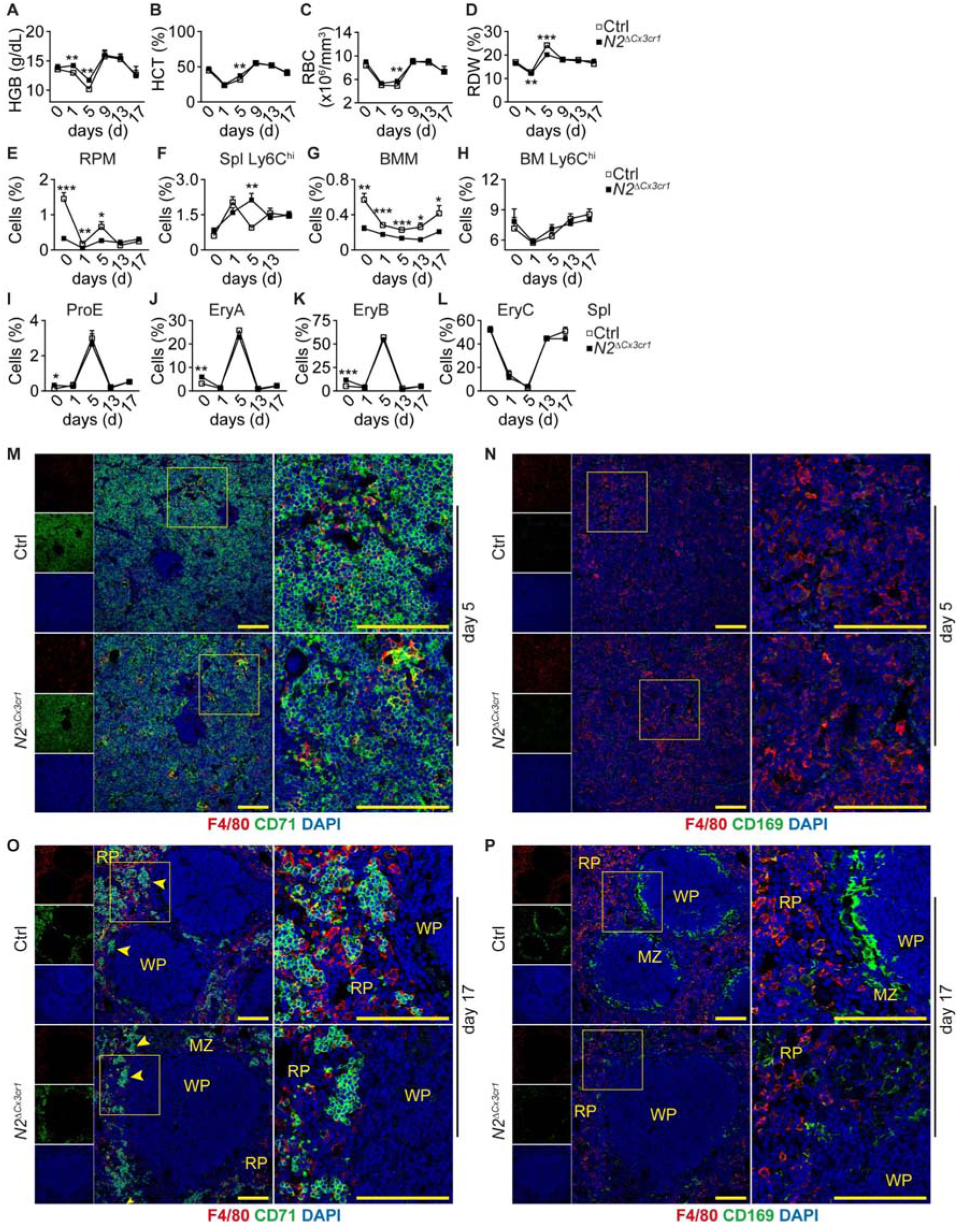
*Notch2*-deficiency confers improved adaptation to PHZ-induced hemolytic anemia. (A-D) Erythroid parameters in Ctrl and *N2*^Δ*Cx3cr1*^ mice injected with PHZ (n = 10-12). (E-H) Relative frequency of (E) RPM, (F) Spl Ly6C^hi^ monocytes, (G) BMM and (H) BM Ly6C^hi^ monocytes in PHZ-treated mice (n = 6-8). (I-L) Relative frequency of Spl erythroid progenitors in PHZ-treated mice (n = 3-8). (M-P) CLSM images of Spl (N, O) on day 5 or (P, Q) 17 after PHZ-injection. Representative of two or more experiments. Arrowheads depict erythropoietic clusters. Scale bar 100 µm. (M,N and O,P) Images are from the same samples but depicted separately for simplicity. (A-L) Data are mean ± SEM pooled from two or more experiments. *P<0.05; **P<0.01; ***P<0.001; (Student’s *t*-test). See also Figure S9 and S10.

Although specialized in heme metabolism, heme overload can cause RPM cell death, for example in severe hemolytic anemia^52^. In Ctrl mice, PHZ administration led to a strong and persistent reduction in RPM numbers, which only transiently and partially recovered (**Figure 7E, Figure S9O**). Concomitantly, the number of Ly6C^hi^ monocytes on day 1 transiently increased in Spl and decreased in BM, preceding an increase of RPM at day 5, suggesting recruitment and active differentiation of monocytes into RPM, as described by Haldar et al.^17^ (**Figure 7E**, **7F and 7H, Figure S9P and S9R)**. This did not occur in *N2*^Δ*Cx3cr1*^ animals, where Ly6C^hi^ monocytes remained elevated in parallel to low numbers of RPM (**Figure 7E and 7F**). Furthermore, BMM frequency in Ctrl mice decreased strongly after PHZ but partially recovered during the late phase. This was in stark contrast to *N2*^Δ*Cx3cr1*^ animals, which showed strongly reduced BMM frequency at baseline and during follow up (**Figure 7G**, **Figure S9Q**). Thus, the defects in the BMM population observed in mice with *Notch2* loss-of-function are pronounced and persistent.

In response to PHZ, there was a strong increase in splenic early erythrocyte progenitors, and a suppression in EryC, which was comparable between Ctrl and mutant mice (**Figure 7I-7L**). Approximately 9 days post PHZ administration, erythroid parameters in PB return to baseline levels (**Figure 7A-7D**). This coincided with decreased number of erythroid progenitors and rise in EryC number in the Spl, BM, PB and Li, likely due to differentiation of progenitors into mature erythrocytes (**Figure 7I-7L, Figure S10**). Despite normalization of erythroid parameters 17 days after PHZ administration, the cellularity in both the Spl and BM continued to exhibit aberrations, where low levels of RPM and BMM persisted in both Ctrl and knockout animals (**Figure 7E and 7G**). This suggests that rapid generation of new RBC has priority over the recovery of ablated RPM and BMM, which occurs at a much slower rate.

This was also reflected in the cellular and spatial changes in the Spl by CLSM. On day 5 post-PHZ administration, Spl lost the characteristic architecture (**Figure 7M and 7N**), and evidence for an active erythropoietic state, seen by the presence of CD71^+^ erythroblastic islands in RP, persisted until day 17 independent of genotype (**Figure 7O-P**). At the late recovery phase, the Spl of Ctrl mice started to return to the normal structure where distinct RP, MZ and WP were visible (**Figure 7O and 7P**). Notably in Ctrl mice, usual CD169^+^ MMM rings and F4/80 RPM were observed, while *N2*^Δ*Cx3cr1*^ Spl showed the typical phenotype with lack of MMM ring and CD169^+^ cells diffused in RP (**Figure 7P**). Taken together, our data suggests that the enhanced erythropoietic activity in the *N2*^Δ*Cx3cr1*^ animals provides an advantage in the initial recovery phase following PHZ-induced anemia. Moreover, an intact Notch2 is essential for the restoration of the typical splenic structure and recovery of the BMM population following hemolysis.

## Discussion

Here, we show that canonical Notch2 signaling is essential for the development of iron-recycling TRM. Mice with loss of *Notch2* and its downstream mediator *Rbpj* displayed defects in RPM and BMM populations and iron overload in the Spl and BM, while prototypic RPM genes required for iron handling were downregulated. This was accompanied by splenic extramedullary hematopoiesis, expansion of CD163-expressing monocytes and CD169^+^ MMM and changes in splenic microarchitecture. Furthermore, defects in RPM and BMM and iron overload were rescued by early postnatal transfer of bone marrow and fetal liver progenitors in a *Notch2*-dependent manner, demonstrating the potential to restore defective TRM niches by Notch-competent progenitors.

Analogous to reports from *Spic-*^18^ and *Pparg*-deficient mice^21^, deletion of *Notch2* led to RPM and BMM reduction, suggesting the possibility that *Spic*, *Pparg* and *Notch2* may operate within a shared or overlapping pathway regulating differentiation and function of these macrophages. It may also indicate that *Notch2* regulates the numbers or behavior of a shared progenitor, which could explain the selective macrophage loss in these specific organs. Although it is widely accepted that the majority of TRM originate from distinct fetal progenitors during embryonic and fetal liver developmental waves^42,53^, evidence for the precise progenitor of RPM and BMM is scarce. In addition, niche-specific signals, derived from the microenvironment, play a crucial role in shaping the identity and function of macrophages, regardless of origin^54^. Defective TRM pools can be reconstituted by transfer of FL or BM cells in newborn mice supporting niche-mediated instruction of macrophage development^21^ ^55^ ^54^. Our reconstitution studies demonstrate that BMM and RPM in *Notch2*-deficient mice are reconstituted by FL or BM cells in a *Notch2*-dependent manner. Several lines of evidence suggest a unique, but possible shared origin and developmental pathway of RPM and BMM, as was suggested by Haldar et al.^17^: (1) the isolated defect of RPM and BMM in *Notch2*-mutant mice not affecting other TRM (such as KC in the liver), (2) the localized iron deposition in their resident niche, (3) the fact that both populations are reconstituted via shared progenitors. The complex spatial and temporal overlaps combined with the lack of appropriate tools make it challenging to precisely determine the timing and extent to which *Notch2* may influence RPM or BMM development. Similarly, the extent to which BM- or FL-derived RPM or BMM are transcriptionally equivalent to their bona fide embryonic-derived counterparts is not addressed in this study and requires further investigation.

Our data also demonstrate that the normal splenic architecture was disrupted in *Notch2*-deficient mice. In the absence of RPM, the MMM ring forming the border between white and red pulp was missing and the expanding pool of CD169^+^ cells accumulated in the RP, partially resembling results from LXRα-deficient (Liver-X Receptor alpha, encoded by *Nr1h3*) mice, which lack splenic MZ macrophages, including MMM^56^. LXRα-deficient mice, however, retain a normal RPM pool with no signs of iron deposits. The structural loss of the CD169^+^ MMM ring in *N2*^Δ*Cx3cr1*^ mice suggests that LXRα signaling might also be regulated via *Notch2*. Indeed, we saw a reduced expression of *Nr1h3* in the absence of *Notch2* using an *in vitro* model mimicking macrophage differentiation. Moreover, reconstitution of *N2*^Δ*Cx3cr1*^ mice with Ctrl BM monocytes rescued the MMM ring defect, similar to BM reconstitution studies in LXRα-deficient mice^56^. Alternatively, loss of RPM may simply allow MMM to expand and translocate into the RP, indicative of either a compensatory mechanism to RPM loss or altered signaling in this macrophage subset due to the lack of *Notch2*. The mechanism and the functional significance of such translocation remain to be explored.

DLL4-Notch-RBPJ signaling induces the expression of LXRα and Spi-C in BM monocytes and enables their differentiation into KC^31,32^. In the spleen, the ligand DLL1 is prominently expressed by endothelial cells in the MZ and regulates splenic MZ B-cell development but also monocyte conversion in a *Notch2*-dependent manner ^57,25^. Indeed, *Dll1* is highly expressed in splenic capillary EC, while *Dll4* expression is limited to capillary arterial EC ^58^. These Dll1-expressing EC could regulate the differentiation of potential precursors into RPM, similar to the process observed for Dll4^31^ and this may be contingent on *Notch2*. In line with these data, our *in vitro* culture experiments show *Notch2*-dependent upregulation of *Nr1h3*, *Spic*, and *Slc40a1* in the presence of DLL1, which was further enhanced by hemin. This suggests that DLL1-Notch2 and Heme-Bach1 pathways closely interact to promote the development of macrophages.

In the spleen, the myeloid growth factor CSF-1 is provided by WT1^+^ RP reticular fibroblasts which is required for RPM maintenance^16^, while the CSF-1 receptor antagonist GW2580 reduced the number of RPM in mice, supporting a role for CSF-1 in RPM survival^36^. IL-33 has also been identified as a regulator of RPM development^20^. While it is evident that Notch, CSF-1-, and IL-33 signaling axes are needed for the formation or maintenance of RPM, the link between these pathways remains to be established.

The neonatal Spl contains CD71-expressing erythroid precursors and is an active site of erythropoiesis but their numbers gradually decrease and in adult mice, the BM becomes the primary site of erythropoiesis^41^. In adult *N2*^Δ*Cx3cr1*^ mice, we found signs of iron overload and persistence of splenic erythroid progenitors, indicative of extramedullary erythropoiesis, which was paralleled by reduced erythroid progenitors and BMM in the BM. These findings are consistent with ineffective or insufficient hematopoiesis in the bone marrow, causing the persistence of extramedullary hematopoiesis as a fully compensatory response, since no PB blood count abnormalities were noted. This may further be promoted by increased splenic iron levels, which enhances the erythropoietic potential by maintaining erythroid progenitors and promoting their differentiation into mature erythrocytes^59^. Furthermore, loss of CD169^+^ BMM facilitates the egress of hematopoietic stem/progenitor cells (HSC) from the BM into circulation^60^, and this mechanism may further explain the changes seen in *N2*^Δ*Cx3cr1*^ mice, i.e. increase in ProE in the Spl and a reciprocal decrease in BM. However, the lack of Ki-67 staining in these CD71^+^ islands might suggest an HSC proliferation tropism to the BM before egress to the spleen. Thus, Notch2-dependent changes in the BMM and RPM likely shift erythropoiesis to the spleen.

We propose that maintenance of high numbers of erythroblasts in the RP offers a protective advantage during acute hemolytic anemia. This may have significant implications for traumatic blood loss, where early mitigation could substantially influence recovery outcomes. Nevertheless, *N2*^Δ*Cx3cr1*^ mice are still able to recover from hemolysis, indicating that other key regulators beyond RPM or BMM are efficiently involved in restoring homeostasis. However, it still remains to be explored whether the loss of RPM and BMM with the associated structural changes might have an influence on iron homeostasis and erythropoiesis in iron deficiency conditions or on responses to infection. A recent study^4^ has directly linked RPM to the clearance of *Streptococcus pneumoniae*, wherein the absence of RPM had detrimental effects on recovery after infection. Given their use of *Spic*-deficient mice to elucidate this, it is plausible that *N2*^Δ*Cx3cr1*^ mice may exhibit somewhat similar responses.

Limitations: Cx3cr1 is broadly expressed throughout the myeloid lineage, both during embryonal development and postnatally. Thus, constitutive *Cx3cr1^Cre^*-based targeting is not able to identify the exact origin of macrophage progenitors. For this aim, new, more precise tools are needed to selectively target specific progenitors at different stages of development.

Our findings reveal a previously unknown critical role of canonical Notch2 signaling as a master regulator of RPM and BMM development. These results highlight the indispensable function of Notch2 for iron-recycling macrophages, iron homeostasis and erythropoiesis, which affects splenic structure.

## Acknowledgments

We thank the Central Animal Facility, Research Core Facility Cell Sorting, Research Core Unit Genomics, and Research Core Unit for Laser Microscopy of Hannover Medical School for their excellent support. Supported by grants from DFG LI 948/10-1 to F.P.L. and GA 2443/3-1 to J.G.

## Author Contributions

F.N.S., S.S., T.K., Y.X. and J.G. performed experiments. F.N.S.: Conceptualization, Data curation, Formal analysis, Validation, Visualization, Methodology, Writing - original draft, review and editing; T.K.: Conceptualization, Data curation, Validation, Methodology, Writing - review and editing. SS and YX: Resources, Data curation; review and editing; B.C and Ti.K.: Methodology; L.D. and A.K.: Methodology; M.L.: Resources, Methodology; H.H. and K.S.O.: Resources, Writing – review; J.G.: Conceptualization, Project administration, Data curation, Funding acquisition, Validation, Methodology, Supervision, Writing - original draft, review and editing; F.P.L.: Conceptualization, Resources, Supervision, Funding acquisition, Validation, Writing - original draft, review and editing.

## Supplemental information

Document S1: Figures S1-S10. and Table S1.

## Methods

### Mice

All mice used are adults (8-16 weeks old) and experiments performed were done with age and sex-matched littermate controls, unless otherwise indicated. CD45.1^+^ mice were from the central animal facility of Hannover Medical School (ZTL, MHH). *Cx3cr1^Cre^* mice^38^, *Cx3cr1^GFP/+^*^44^, *Notch2^lox/lox^* mice^39^, *Rbpj^lox/lox^* ^40^, *ROSA26-mT/mG* mice^48^ have been previously described. *N2*^Δ*Cx3cr1*^ or *Rbpj*^Δ*Cx3cr1*^ mice were generated by crossing *Cx3cr1^Cre^* mice^38^ with *Notch2^lox/lox^*^39^ or *Rbpj^lox/lox^* animals^40^. *N2*^Δ*Cx3cr1*^ *mG* and Ctrl *mG* strains were generated by crossing *ROSA26-mT/mG* mice^48^ with *Cx3cr1^Cre^ Notch2^lox/lox^ (N2*^Δ*Cx3cr1*^) or *Cx3cr1^Cre^ Notch2^+/+^*(Ctrl) animals, respectively. All experiments performed were approved by the local animal welfare board (LAVES, Lower Saxony). Mice were bred and housed under specific pathogen-free conditions in the central animal facility of Hannover Medical School (ZTL, MHH).

### Tissue and Cell Preparation

To prepare single-cell suspensions, mice were euthanized and organs were collected as previously described^37^. Briefly, Spl were mechanically dissociated, BM was collected by centrifugation and PB was collected from the inferior vena cava. Livers were digested at 37°C for approximately 1 h in DMEM medium supplemented with 500 U/ml Collagenase II (Worthington). Erythrocytes were excluded using red blood cell lysis buffer (BioLegend) or by density centrifugation using Histopaque 1083 (Sigma-Aldrich). Cells were thoroughly washed and resuspended in PBS containing 10% fetal calf serum (FCS) and 2 mM EDTA and used for flow cytometry. For splenic macrophage subset analysis, spleens were digested on ice for 30 min. in DMEM medium supplemented with 500 U/ml Collagenase II (Worthington), 1.5 U/ml DNAse and 2% FCS.

### Flow Cytometry and Cell Sorting

To reduce non-specific binding of antibodies to Fc-receptors, blocking with anti-mouse CD16.2 (FcγRIV) and anti-mouse CD16/32 (TruStain FcX™) in single-cell suspensions prepared from Spl, PB, BM or Liver was performed. Cells were then thoroughly washed, labeled with primary and secondary antibodies or streptavidin-fluorochrome conjugates and used for flow cytometry (LSR II, BD Biosciences) or for sorting (FACSAria III Fusion, FACSAria IIu, BD Biosciences). All flow cytometry data were analyzed using FlowJo software (FlowJo LLC).

FSC and SSC traits were used to identify the cells. After doublet exclusion (based on SSC-W and SSC-A traits), the relative frequency or absolute numbers of each subset was determined (calculated from live cell gate, normalized per Spl, per mg BM or per µl PB). Unsupervised t-distributed stochastic neighbor embedding (t-SNE) analysis^61^ was performed on live CD45^+^Lin^neg^CD117^neg^ population after exclusion of CD11b/CD11c double negative cells in concatenated samples using FlowJo software.

### Apoptosis and Proliferation Assay

Apoptosis assay was performed according to the manufacturer’s instructions (Biolegend). Briefly, single-cell suspensions were stained with primary and secondary antibodies or streptavidin-fluorochrome conjugates, washed and resuspended in AnnexinV (AnnV) binding buffer (Biolegend). Cells were stained with AnnV at room temperature for 20 min. After incubation, propidium iodide (PI) (Sigma-Aldrich) was added and cells were immediately analyzed by flow cytometry.

BrdU incorporation assay was performed according to the manufacturer’s instructions (BD Pharmingen). Briefly, 30 µl BrdU (10 mg/ml in sterile PBS) was administered to neonates intraperitoneally (P5-P7). The animals were sacrificed the next day. Single cell suspensions were stained with surface antibodies, fixed-permeabilized and stained with anti-BrdU. BrdU incorporation was analyzed using flow cytometry. Antibodies used are described in Key Resources Table.

### Immunohistochemistry and Microscopy

Organs were harvested from euthanized mice, fixed in 4% PFA in PBS and embedded in optimum cutting temperature (OCT) compound. 8 µm thick cryosections were prepared using a cryostat (Leica CM3050S) and stored at -20°C. For immunostaining, sections were blocked with anti-mouse CD16/CD32 (TruStain fcX from BioLegend) in 3% BSA and stained with appropriate primary and secondary antibodies. Sections were counterstained with DAPI. Images were taken using a Leica Inverted 3 TCS SP8 DMi8 and images were processed with Leica Application Suite (LAS) AF Lite software (Leica).

### Prussian Blue Iron Staining

Ferric iron deposits were visualized using Perl’s Prussian blue stain in frozen tissue sections and counterstained with nuclear red. Multiple images of each biological replicate were taken using 10x and 20x objectives of Leica DMI 3000B DFC420C microscope. Images were acquired and processed using LAS v3 (Leica).

### Iron Quantification

Unless otherwise stated, stocks were prepared by dissolving solutes in water. The solutions were prepared as follows: Homogenizing buffer (HB) was prepared by dissolving sodium chloride (Roth) in NaOH-HEPES (pH 7.4, Sigma-Aldrich) at final concentrations of 0.15 M and 10 mM, respectively. Extraction buffer (EB) was prepared by combining 50% trichloroacetic acid (Sigma-Aldrich) and 8% sodium pyrophosphate tetrabasic (Sigma-Aldrich) and kept at 4°C. Ferrozine solution (FS) was freshly prepared before each use by combining 38 mM ascorbic acid (Sigma-Aldrich), 1.3 mM ferrozine (Sigma-Aldrich) and 1.4 M (pH 4.8) of sodium acetate (Sigma-Aldrich). For Fe standards, a stock concentration of 1 mM FeCl_3_ (Sigma-Aldrich) was prepared in 10 mM HCl (Sigma-Aldrich). Standard curves were generated at a FeCl_3_ starting concentration of 100 µM.

Organs were manually disrupted/dissolved in 300 µl HB followed by addition of EB (1:1 ratio of HB and EB). This was digested at 95°C for 20 min. followed by centrifugation at 13,000 rpm for 10 min. 25 µl of the resulting supernatant was mixed with 75 µl FS in 96 half well flat bottom plates (Greiner) and absorbance was measured at 562 nm (Infinite® 200, Tecan). Each sample was analyzed in duplicates and the average value was used for calculation of Fe concentration per mg organ or per μl blood.

### Adoptive Transfer to Neonates

BM or FL cells were harvested from 8-10 weeks old or E14.5 embryo donor mice, respectively. Single cell suspension was prepared in PBS. 8×10^6^ donor cells were injected once intraperitoneally to neonates (P5-P7). In a separate experiment, BM Ly6C^hi^ monocytes were sorted from adult mice and used for reconstitution experiment. Recipient mice were euthanized 8-14 weeks later and reconstitution was analyzed using flow cytometry and confocal microscopy.

### *In vitro* conversion Studies

24-well plates were coated at room temperature for 2-3 h with IgG-Fc or DLL1-Fc ligands (all from R and D) reconstituted in PBS. Sorted BM Ly6C^hi^ monocytes were cultured for four days in coated plates in the presence of CSF-1 (10ng/ml, Peprotech) at 37°C. Cells were stimulated on day 2 with 40 µM porcine Hemin (Sigma-Aldrich) reconstituted in 2 mM NaOH (Sigma). Cultured cells were harvested and isolated RNA was used for gene expression analysis (QRT-PCR).

### RNA Isolation and Quantitative real-time PCR

Purification of total RNA from cell lysates was done using the Nucleospin™ RNA plus kit (Macherey-Nagel™) and reverse transcribed using SuperScript™ III First-Strand Synthesis System (Invitrogen) according to the manufacturer’s instructions. The resulting cDNA was used for quantitative real-time PCR (QRT-PCR) using specific primers and FastStart Essential DNA Green Master on a LightCycler 96 system (Roche). The resulting gene expresssion was normalized to RPS9 and values were calculated using the comparative CT method (2-DDCT)^62^. Primer sequences are as follows:

*Spic* forward: 5’-CCA CTT GGT TTT CCT GAA CGT-3’

*Spic* reverse: 5’-TTG CGG AAA TGT CAG CGA GTA-3’

*Hmox1* forward: 5’-GCC GAG AAT GCT GAG TTC ATG-3’

*Hmox1* reverse: 5’-TGG TAC AAG GAA GCC ATC ACC-3’

*Nr1h3* forward: 5’-TGT GCG CTC AGC TCT TGT-3’

*Nr1h3* reverse: 5’-TGG AGC CCT GGA CAT TAC C-3’

*Slc40a1* forward: 5’-CTA CCA TTA GAA GGA TTG ACC AGC T-3’

*Slc40a1* reverse: 5’-CAA ATG TCA TAA TCT GGC CGA-3’

*Pparg* forward: 5’-GAA AGA CAA CGG ACA AAT CAC C-3’

*Pparg* reverse: 5’-GGG GGT GAT ATG TTT GAA CTT G-3’

*Cd163* forward: 5′-CAT GTG GGT AGA TCG TGT GC-3’

*Cd163* reverse: 5′-TGT ATG CCC TTC CTG GAG TC-3’

*RPS9* forward: 5’-GGA TTT CTT GGA GAG GCG GC-3’

*RPS9* reverse: 5’-ACC TGC TTG CGG ACC CTA AT-3’

### Phenylhydrazine treatment

Mice were administered 2 mg/mouse phenylhydrazine (PHZ, Sigma-Aldrich) in PBS. Mice were sacrificed at different time points after anemia induction and organs of interest were collected for analysis. For blood parameter analysis, blood was drawn from the cheek and measured on a hematology analyzer (Scil Vet ABC, Antech Company, Viernheim, Germany).

### Statistical analyses

All statistical analyses were performed using Graphpad Prism. Results are shown as mean ± standard error of mean (SEM). N numbers represent biological replicates and all experiments were performed at least two times unless otherwise stated. Groups were compared using unpaired two-tailed Student’s t-test with confidence interval of 95%. For comparison of multiple experimental groups, one-way or two-way analysis of variance (ANOVA) with Bonferroni’s multiple comparison post test was used.

