## Supplementary Figures and Table for "Canonical Notch2 Signaling Regulates the Development of Iron-recycling Macrophages and Iron Homeostasis"

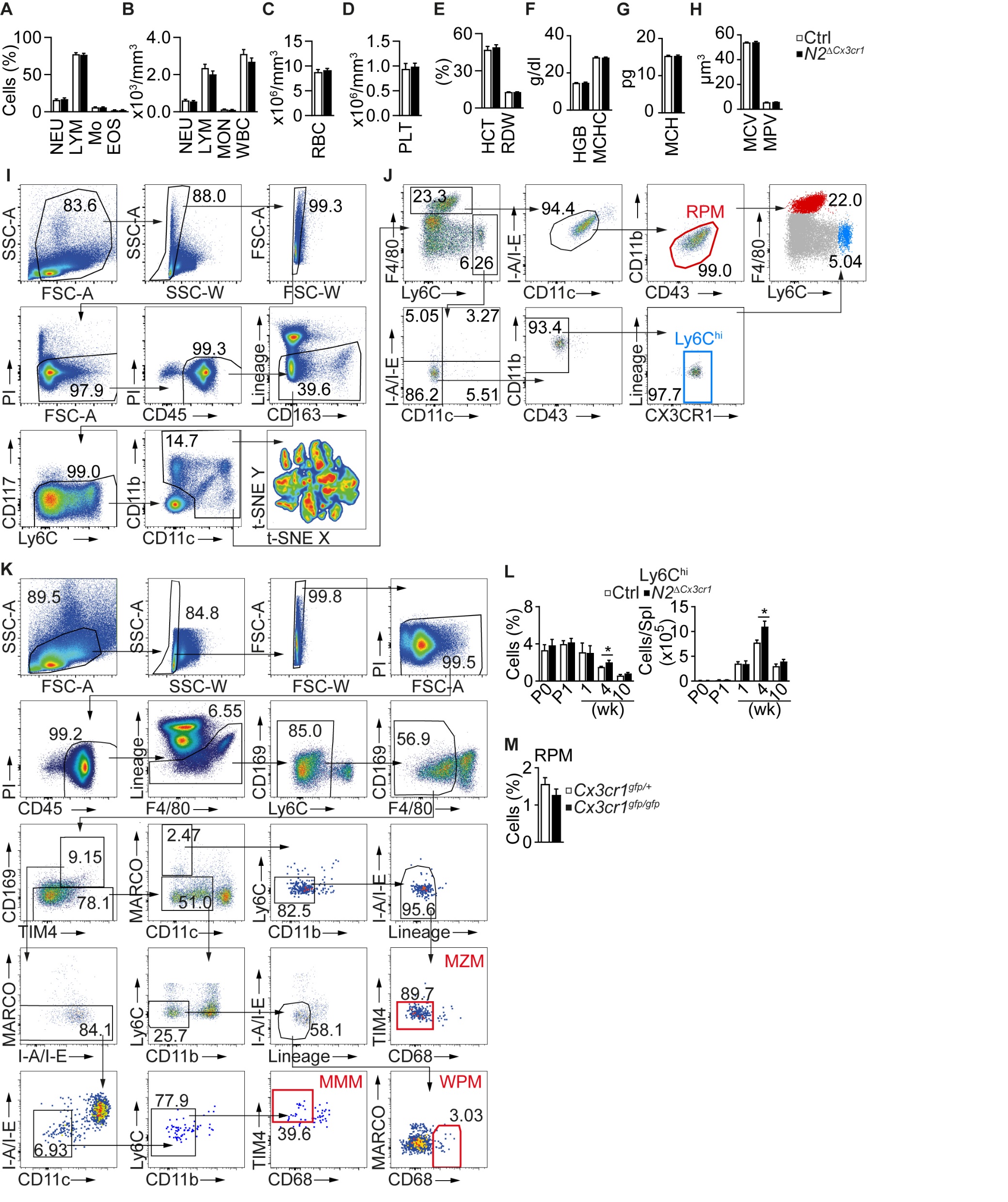

**Figure S1. Related to Figure 1. Gating strategies and general characterization of MPS.**

(A-H) Complete blood counts from 10 wk old Ctrl or *N2^ΔCx3cr1^* mice. (n=6-8)

(I) Gating strategy for t-SNE analysis and for definition of splenic phagocytic cells. Doublets, dead (PI^+^), CD45^neg^, Lin^+^, CD117^+^ and CD11b/CD11c-double-negative cells were excluded from Ctrl or *N2^ΔCx3cr1^* spleen samples prior to down-sampling, concatenation and t-SNE analysis.

(J) Gating strategy for definition of splenic RPM and Ly6C^hi^ monocytes for conventional flow cytometry analysis. BMM and BM Ly6C^hi^ monocytes were defined using the same gating strategy for splenic RPM and Ly6C^hi^ monocytes.

(K) Gating strategy for the definition of other splenic macrophage populations for conventional flow cytometry analysis.

(L) Relative and (M) absolute frequency of splenic Ly6C^hi^ monocytes in Ctrl or *N2^ΔCx3cr1^* mice (n=3-8).

(M) Bar graphs showing relative frequency of RPM populations in *Cx3cr1^gfp/+^* or in *Cx3cr1^gfp/gfp^* mice (n=6-8).

(A-H, L,M) Data are mean ± SEM and pooled from, or representative of at least two independent experiments. *P<0.05; **P<0.01; ***P<0.001; (Student’s *t*-test). Lin: Lineage (CD3, CD19, B220, Ly6G, Terr119, NK1.1).

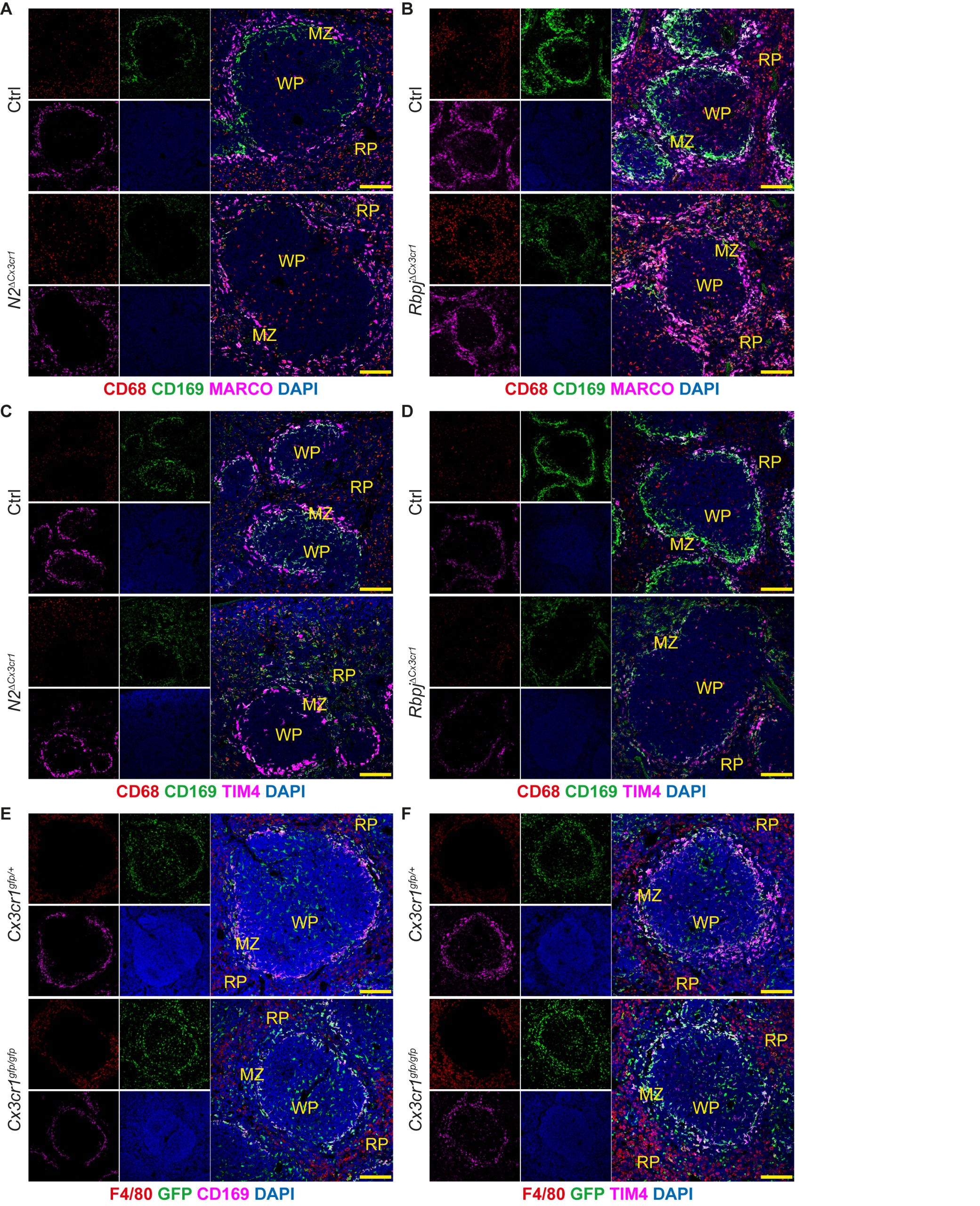

**Figure S2. Related to Figure 3. Impaired splenic microarchitecture and iron homeostasis in canonical *Notch2* signaling-deficient mice**

(A-D) CLSM images of Spl from 10 wk old (A,C) Ctrl or *N2^ΔCx3cr1^* or (B,D) Ctrl or *Rbpj^ΔCx3cr1^* mice.

(E,F) CLSM images of Spl from 10 wk old (A,C) *Cx3cr1^1gfp/+^* or *Cx3cr1^1gfp/gfp^* mice.

(A-F) Data are representative of at least two or more independent experiments. Scale bar 100 µm.

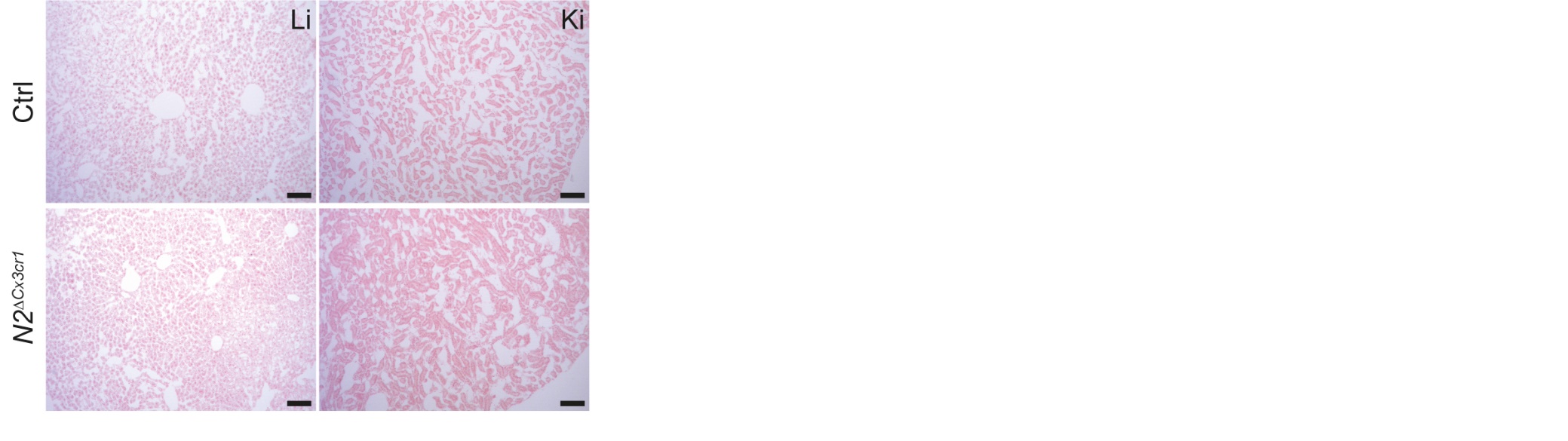

**Figure S3. Related to Figure 3. Absence of iron deposits in the Liver and Kidney**

Perl’s Prussian blue staining of Liver (Li) and Kidney (Ki) of 10 wk old Ctrl or *N2^ΔCx3cr1^* mice. Representative microscopy images from at least two experiments. Scale bar 100 µm.

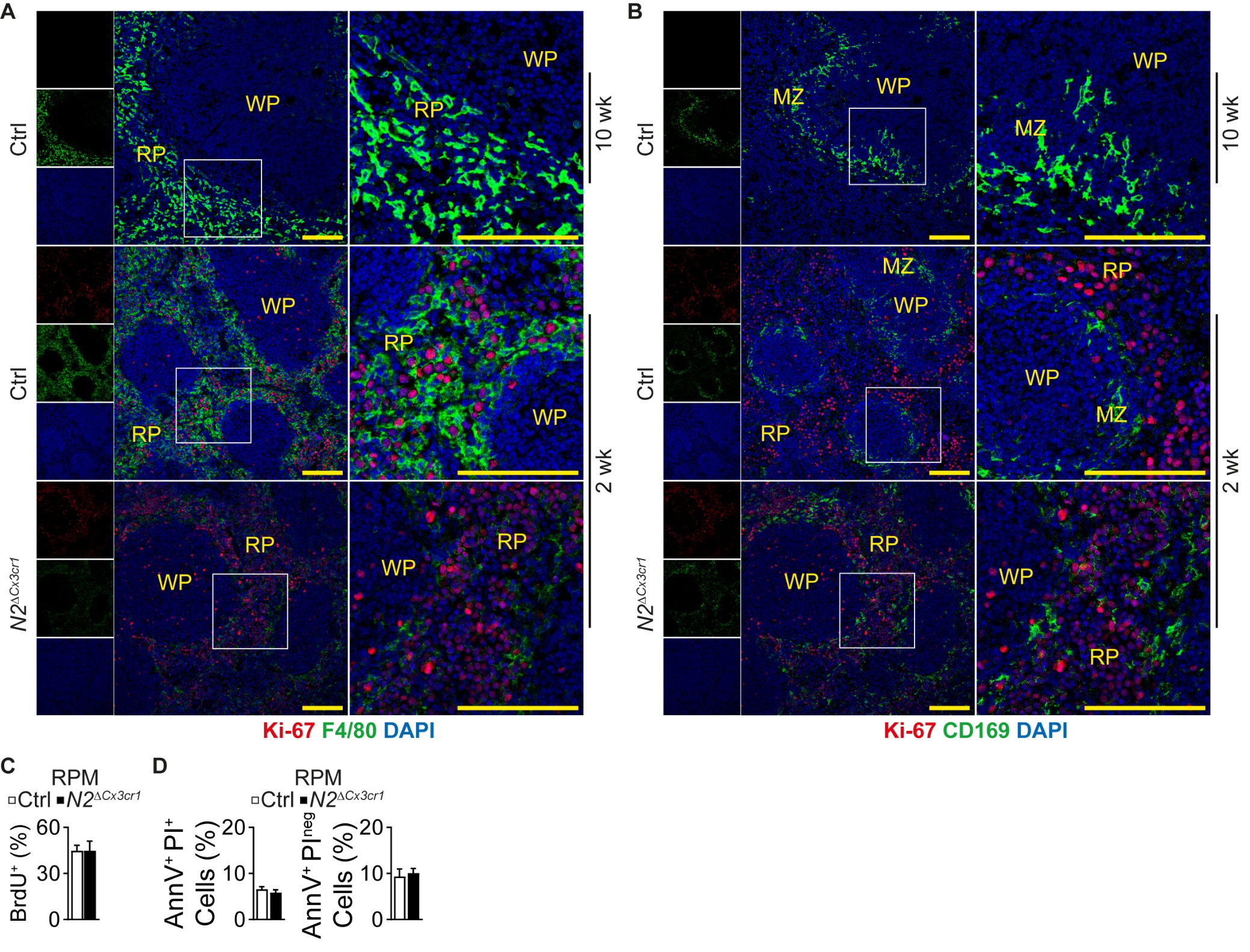

**Figure S4. RPM Proliferation and Apoptosis.**

(A-B) CLSM images of Spl from 10 wk- or 2 wk-old Ctrl or *N2^ΔCx3cr1^* mice. Images are derived from the same sample but depicted separately for simplicity. Representative of two experiments. Scale bar 100 µm.

(C) Frequency of BrdU^+^ RPM in 2 wk old Spl (n=3-5)

(D) Frequency of AnnV^+^PI^+^ or AnnV^+^ PI^neg^ RPM in 10 wk old mice (n=3-4).(C-D) Data are mean ± SEM and pooled from, or representative of at least two independent experiments. *P<0.05; **P<0.01; ***P<0.001; (Student’s *t*-test).

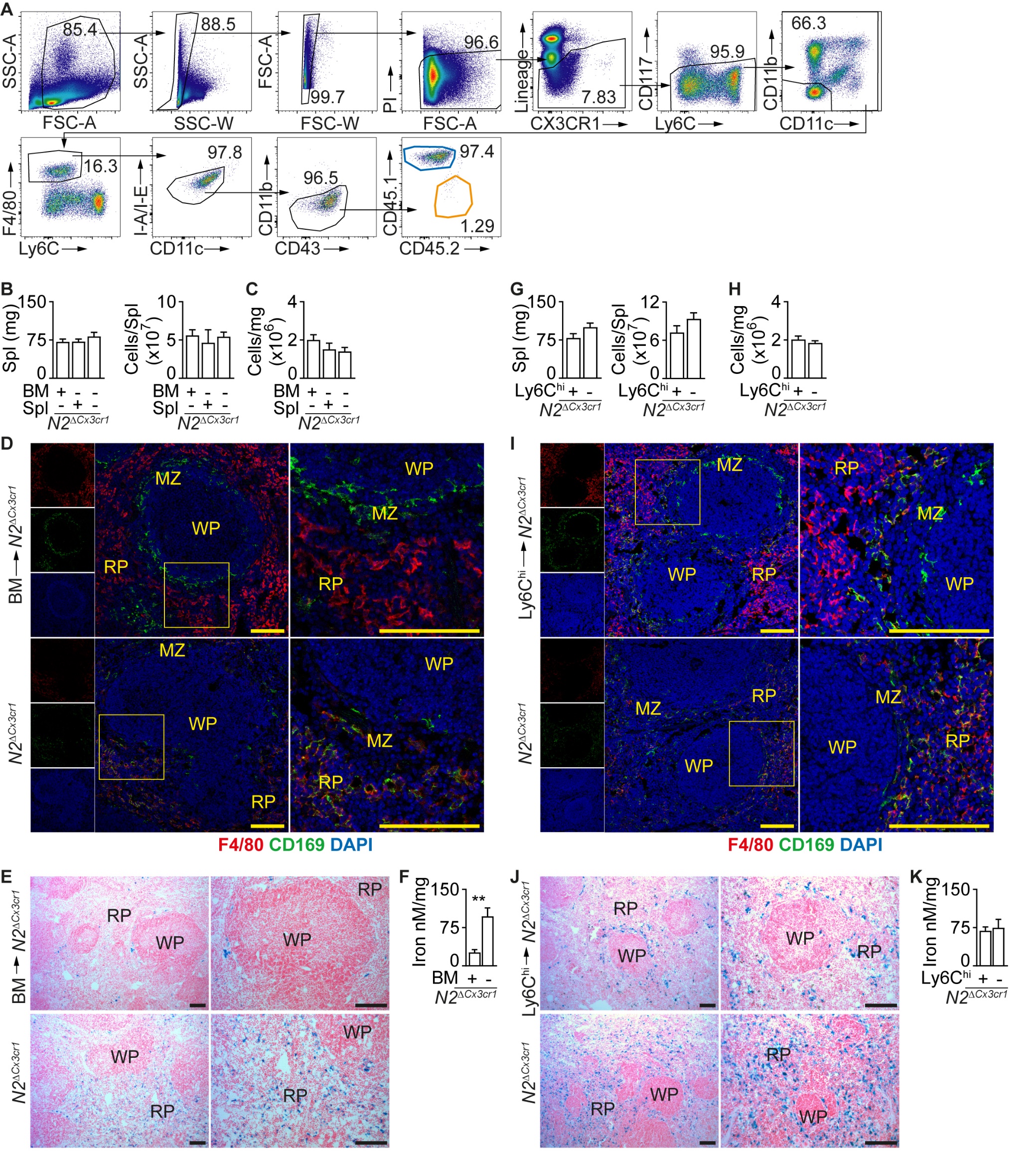

**Figure S5. Related to Figure 4. Reconstitution of the splenic niche with BM monocyte-derived RPM**

(A) Complete gating strategy for identification of congenic CD45.1^+^ donor cells in recipient *N2^ΔCx3cr1^* mice.

(B) Spl weight and total cell number of mice reconstituted with BM or Spl cells.

(C) Frequency of recipient BM cells normalized per mg BM after reconstitution with BM or Spl cells.

(D) CLSM images of Spl from BM-reconstituted or non-reconstituted recipient *N2^ΔCx3cr1^* mice.

(E) Perl’s Prussian blue staining showing iron deposits in Spl of BM-reconstituted or non-reconstituted recipient mice. Representative of at least two experiments. Scale bar 100 µm.

(F) Iron quantification in Spl of BM-reconstituted or non-reconstituted *N2^ΔCx3cr1^* mice.

(G) Spl weight and total cell number of *N2^ΔCx3cr1^* mice after reconstitution with BM Ly6C^hi^ monocytes.

(H) Frequency of BM cells in recipient mice normalized per mg BM after reconstitution with BM Ly6C^hi^ monocytes.

(I) CLSM images of Spl from BM Ly6C^hi^ monocyte-reconstituted or non-reconstituted recipient *N2^ΔCx3cr1^* mice.

(J) Perl’s Prussian blue staining showing iron deposits in recipient Spl of BM Ly6C^hi^ monocyte-reconstituted or non-reconstituted recipient mice. Representative of at least two experiments. Scale bar 100 µm.

(K) Iron quantification in Spl of BM Ly6C^hi^ monocyte-reconstituted or non-reconstituted *N2^ΔCx3cr1^* mice.

(B,C,F-H, K) Data are mean ± SEM and pooled from, or representative of at least two independent experiments (n= 3-6). (B,C) *P<0.05; **P<0.01; ***P<0.001; Ordinary one way ANOVA with Bonferroni’s multiple comparison test. (F-H,K) *P<0.05; **P<0.01; ***P<0.001; (Student’s *t*-test).

(D,I) Images are derived from the same samples as in (D) Figure 4G, 4H and (I) 4O,4P respectively. Depicted as separate images for simplicity. Representative of at least two experiments. Scale bar 100 µm.

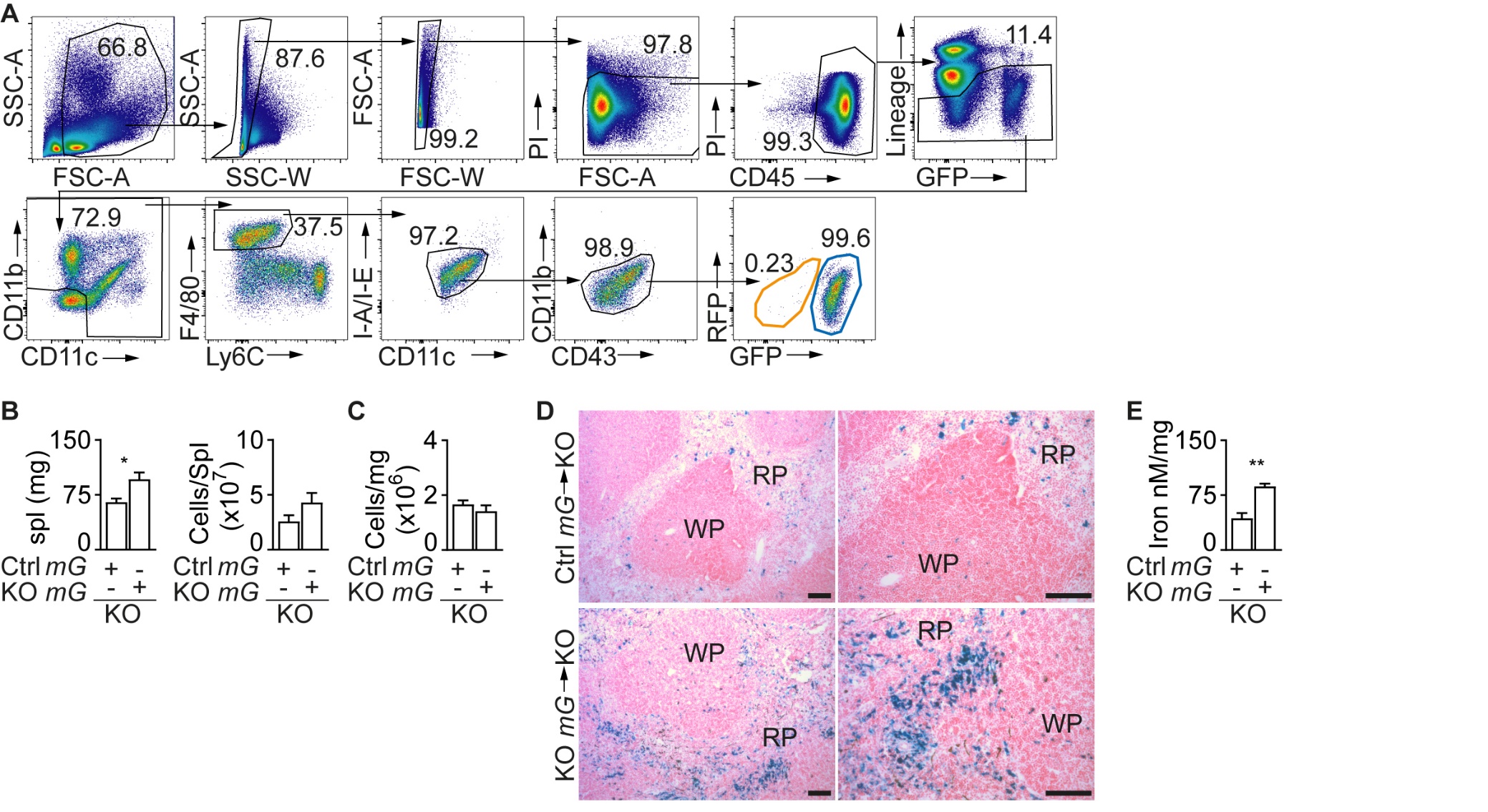

**Figure S6. Related to Figure 5. Development of RPM and BMM requires intrinsic Notch2 signaling**

(A) Complete gating strategy for identification of donor cells in recipient *N2^ΔCx3cr1^* mice.

(B) Spl weight and total cell number in recipient mice reconstituted with Ctrl *mG* or KO *mG* BM.

(C) Frequency of recipient BM cells normalized per mg after reconstitution with Ctrl *mG* or KO *mG* BM.

(D) Perl’s Prussian blue staining showing iron deposits in recipient Spl after reconstitution with Ctrl *mG-* or KO *mG* BM. Representative of at least two experiments. Scale bar 100 µm.

(E) Iron quantification in recipient Spl after reconstitution with Ctrl *mG* or KO *mG* BM.

(B,C,E) Data are mean ± SEM and pooled from two or more experiments (n= 6). *P<0.05; **P<0.01; ***P<0.001; (Student’s *t*-test).

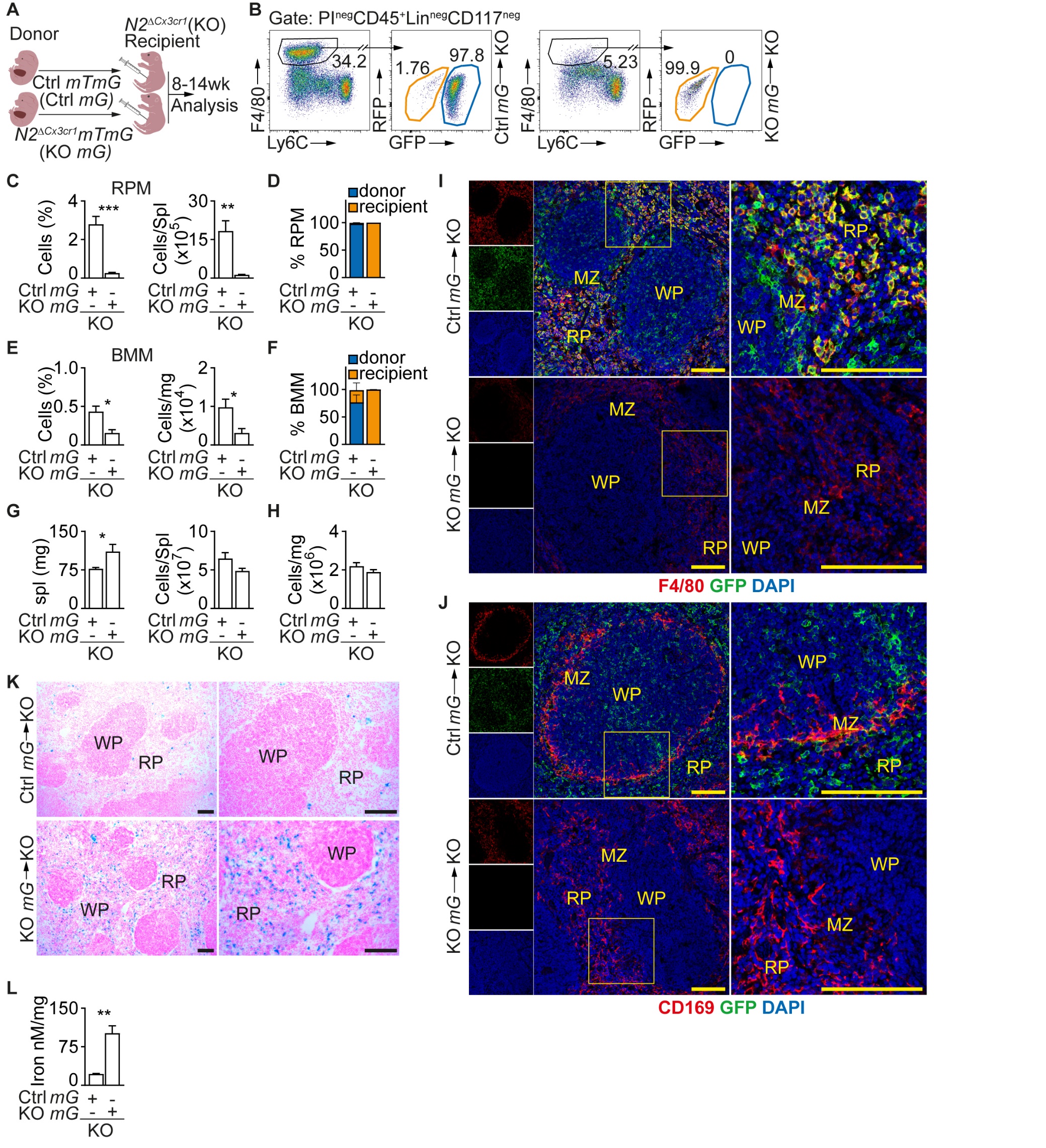

**Figure S7. FL cells give rise to RPM in *Notch2*-dependent manner**

(A) Experimental scheme of reconstitution study using FL cells.

(B) Representative flow cytometry plot depicting donor FL-derived RPM in *N2^ΔCx3cr1^* recipients. Full gating strategy in Figure S6A.

(C) Relative and absolute frequency of RPM in recipient mice.

(D) Frequencies of donor- and recipient-derived RPM in whole RPM pool.

(E) Relative and absolute frequency of BMM calculated per mg BM in recipient mice

(F) Frequencies of donor- and recipient-derived BMM in whole BMM pool.

(G) Spl weight and total cell number in recipient mice reconstituted with Ctrl *mG* or KO *mG* FL cells.

(H) Frequency of recipient BM cells normalized per mg after reconstitution with Ctrl *mG* or KO *mG* FL cells.

(I,J) CLSM images of Spl from Ctrl *mG* or KO *mG* FL-reconstituted *N2^ΔCx3cr1^* mice.

(K) Perl’s Prussian blue staining showing iron deposits in recipient Spl after reconstitution with Ctrl *mG-* or KO *mG* FL cells.

(L) Iron quantification in Spl of Ctrl *mG* or KO *mG* FL-reconstituted *N2^ΔCx3cr1^* mice.

(C-H,L) Data are mean ± SEM and pooled from two experiments (n = 5-6). *P<0.05; **P<0.01; ***P<0.001; (Student’s *t*-test).

(I-K) Representative of at least two or more experiments. Scale bar 100 µm.

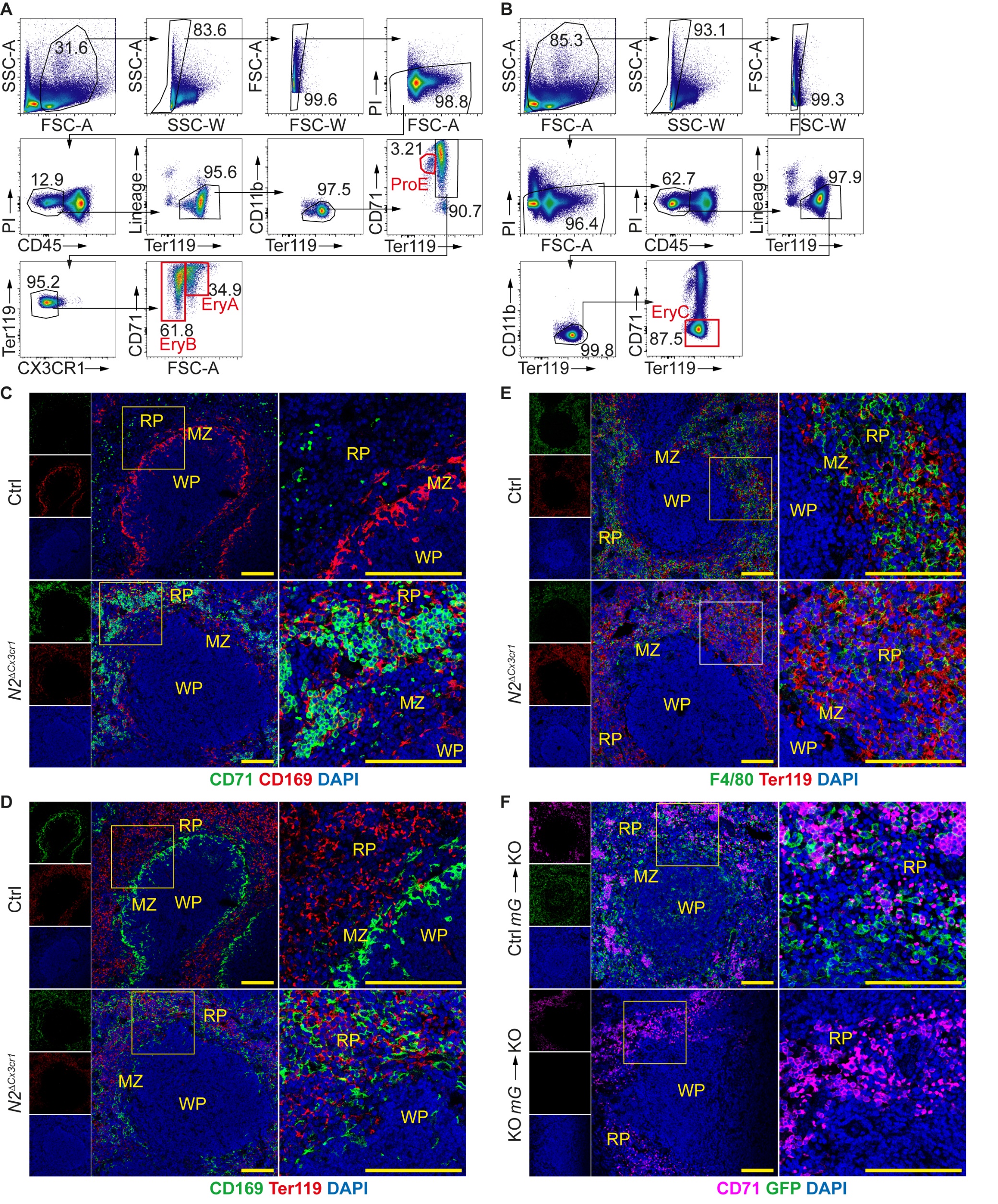

**Figure S8. Related to Figure 6. Enhanced extramedullary erythropoiesis in *N2^∆Cx3cr1^* mice**

(A,B) Complete gating strategy to identify (A) erthyroid progenitors (ProE, EryA, EryB) and (B) mature erythrocytes (EryC). Lin: Lineage (CD3, CD19, B220, Ly6G, CD41, NK1.1). PI: Propidium iodide.

(C-E) CLSM images of Spl from 10 wk old Ctrl or *N2^ΔCx3cr1^* mice. (C,D) Images are derived from the same sample but are shown separately for simplicity. (E) Derived from the same sample as in Figure 6I and 6J but shown separately for simplicity.

(F) CLSM images of Spl from 10 wk old *N2^ΔCx3cr1^* reconstituted with Ctrl *mG* or KO *mG* FL cells.

(C-F) Representative of two or more experiments. Scale bar 100 µm.

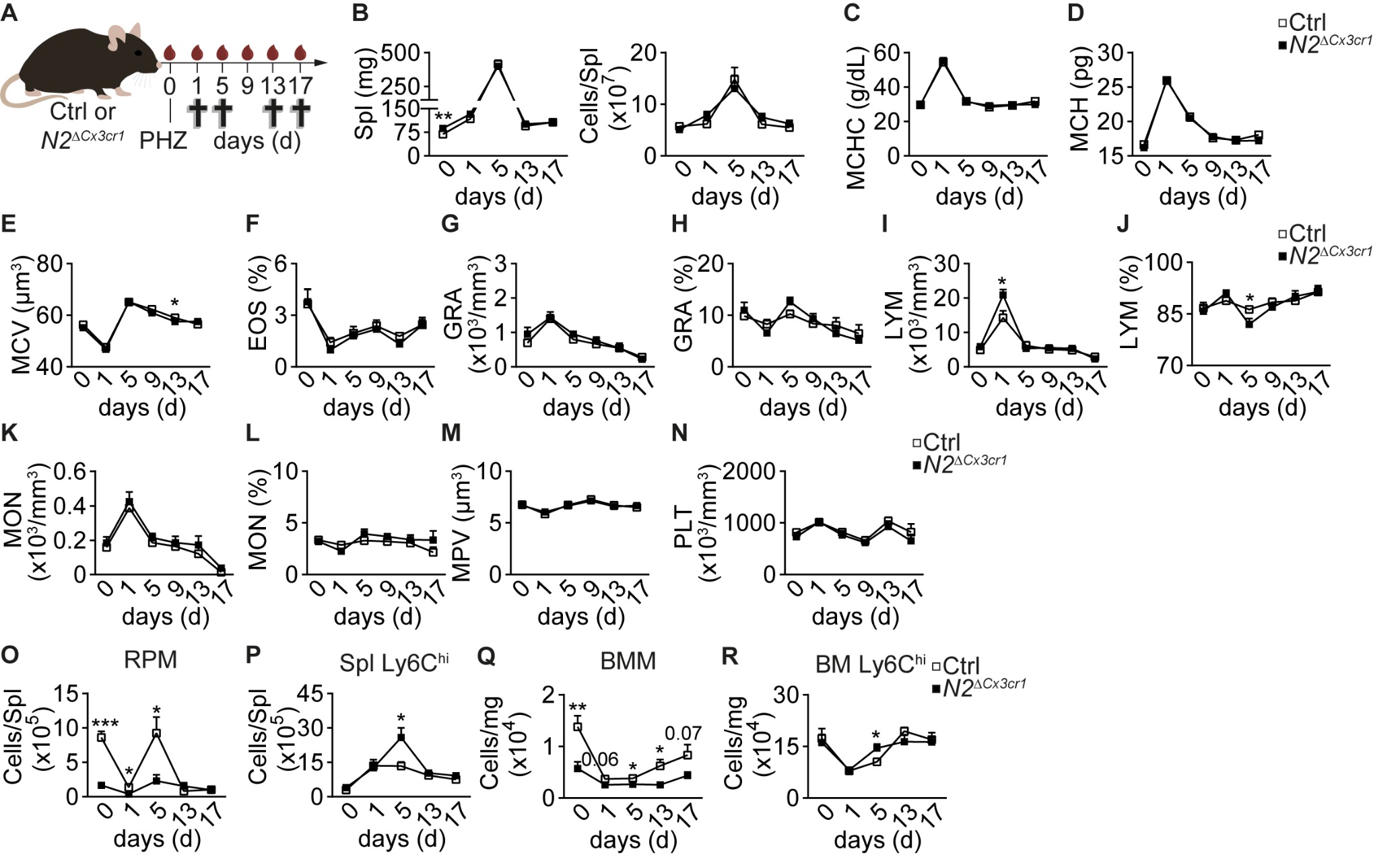

**Figure S9. Related to Figure 7. Notch2-deficiency confers improved adaptation to PHZ-induced hemolytic anemia**

(A) Experimental scheme. Blood samples were collected on indicated days, crosses represent experimental end points.

(B) Spl weight and total cell number after induction of anemia(n=6-8).

(C-N) Erythroid parameters in Ctrl and *N2^ΔCx3cr1^* mice injected with PHZ (n=10-12).

(O,P) Absolute frequency of (O) RPM and (P) Spl Ly6C^hi^ monocytes in PHZ-treated mice (n=6-8).

(Q,R) Frequency of (Q) BMM and (R) BM Ly6C^hi^ monocytes (R) normalized per mg BM in PHZ-treated mice (n=6-8).

(B-R). Data are mean ± SEM pooled from two or more experiments. *P<0.05; **P<0.01; ***P<0.001; (Student’s *t*-test).

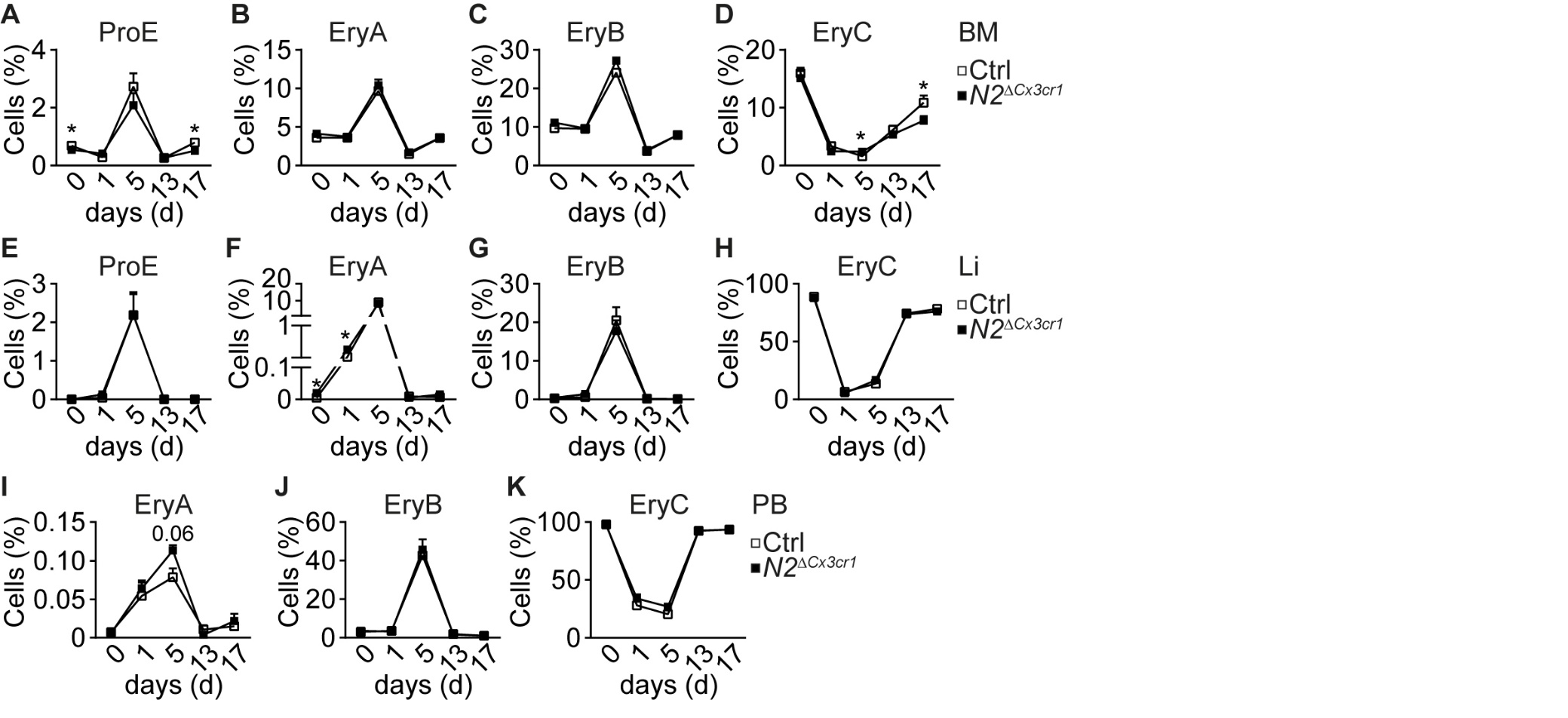

**Figure S10. Related to Figure 7. Erythroid progenitors in PHZ-treated mice.**

(A-K) Relative frequencies of erythroid progenitors in BM (A-D), Liver (Li) (E-H) and PB (I-K) of PHZ-treated mice (n=3-8). Data are mean ± SEM pooled from two or more experiments. *P<0.05; **P<0.01; ***P<0.001; (Student’s *t*-test).

**Table 1: Surface phenotype signatures used for identification of distinct cell populations *in vivo*. Related to Figure 1 and Figure 6.**

| **Population** | **Phenotype** |
| --- | --- |
| RPM/BMM | Live(PI^neg^)CD45^+^Lin^lo/-^CD117^neg^F4/80^hi^Ly6C^lo/-^I-A/I-E^lo/-^CD11c^lo/-^CD11b^lo/-^CD43^lo/-^ |
| MZM | Live(PI^neg^)CD45^+^Lin^lo/-^Ly6C^neg^F4/80^lo^CD169^neg^MARCO^+^CD11b^neg^I-A/I-E^neg^  TIM4 ^neg^CD68^neg^ |
| MMM | Live(PI^neg^)CD45^+^Lin^lo/-^Ly6C^neg^F4/80^lo^CD169^+^MARCO^neg^CD11b^neg^CD11c^neg^  I-A/I-E^lo/-^TIM4^+^CD68^neg^ |
| WPM | Live(PI^neg^)CD45^+^Lin^lo/-^Ly6C^neg^F4/80^lo^CD169^neg^MARCO^neg^CD11b^neg^I-A/I-E^lo/-^CD68^+^ |
| Ly6C^hi^ monocytes | Live(PI^neg^)CD45^+^Lin^lo/-^CD117^neg^F4/80^lo/-^Ly6C^hi^I-A/I-E^neg^CD11c^neg^CD11b^+^  CD43^neg^CX3CR1^+^ |
| ProE | Live(PI^neg^)CD45^neg^Lin^lo/-^CD11b^lo/-^CD71^int^Ter119^int^ |
| EryA | Live(PI^neg^)CD45^neg^Lin^lo/-^CD11b^lo/-^CD71^hi^Ter119^hi^FSC-A^hi^ |
| EryB | Live(PI^neg^)CD45^neg^Lin^lo/-^CD11b^lo/-^CD71^hi^Ter119^hi^FSC-A^lo^ |
| EryC | Live(PI^neg^)CD45^neg^Lin^lo/-^CD11b^lo/-^CD71^lo/-^Ter119^hi^ |

For RPM/BMM, MZM, MMM, WPM, Ly6C^hi^;

**Lin:** CD3, CD19, B220, Ly6G, Ter119, NK1.1

For ProE, EryA, EryB, EryC;

**Lin:** CD3, CD19, B220, Ly6G, CD41, NK1.1
